# Small-molecule targeting of MUSASHI RNA-binding activity in acute myeloid leukemia

**DOI:** 10.1101/321174

**Authors:** Gerard Minuesa, Steven K. Albanese, Arthur Chow, Alexandra Schurer, Sun-Mi Park, Christina Z. Rotsides, James Taggart, Andrea Rizzi, Levi N. Naden, Timothy Chou, Saroj Gourkanti, Daniel Cappel, Maria C. Passarelli, Lauren Fairchild, Carolina Adura, J Fraser Glickman, Jessica Schulman, Christopher Famulare, Minal Patel, Joseph K. Eibl, Gregory M. Ross, Derek Tan, Christina Leslie, Thijs Beuming, Yehuda Goldgur, John D. Chodera, Michael G. Kharas

## Abstract

The MUSASHI family of RNA binding proteins (MSI1 and MSI2) contribute to a wide spectrum of cancers including acute myeloid leukemia. We found that the small molecule Ro 08–2750 (Ro) directly binds to MSI2 and competes for its RNA binding in biochemical assays. Ro treatment in mouse and human myeloid leukemia cells resulted in an increase in differentiation and apoptosis, inhibition of known MSI-targets, and a shared global gene expression signature similar to shRNA depletion of MSI2. Ro demonstrated in vivo inhibition of c-MYC and reduced disease burden in a murine AML leukemia model. Thus, we have identified a small molecule that targets MSI’s oncogenic activity. Our study provides a framework for targeting RNA binding proteins in cancer.

## INTRODUCTION

RNA-binding proteins (RBPs) play critical roles in cell homeostasis by controlling gene expression post-transcriptionally. Ribonucleoprotein complexes are essential for all steps of mRNA processing including splicing, polyadenylation, localization, stability, export and translation^1^. The contribution of RBPs to tumorigenesis (e.g. SRSF2, SF3B1, MSI and SYNCRIP), through genetic perturbation or epigenetic dysregulation, has been found in a variety of human cancers^2–9^. Deregulation of the MSI family of RBPs was initially reported in gliomas^10^, medulloblastomas^11^ and hepatomas^12^. Since then, studies in a diverse range of neoplasms have involved MSI family, including aggressive forms of colorectal^13,14^, breast^15,16^, lung^17^, glioblastoma^18^ and pancreatic cancers^19,20^ and hematological malignancies. Among these tissues, the hematopoietic system has been the most well characterized to dissect MSI function. The *MSI2* gene was initially reported as a translocation partner with HOXA9 in patients progressing from chronic myelogenous leukemia to blast crisis (CML-BC)^21^. More recently, other rare genetic alterations (involving *MSI2, EVI1, TTC40 and PAX5* genes) have been identified in leukemia patients^22,23,24^. *MSI2* expression is detected in 70% of AML patients and it correlates with a poor clinical prognosis in multiple hematological malignancies^25–29^. Thus, MSI2 has been proposed as a putative biomarker for diagnosis as well as a potential therapeutic target for AML^29^.

The relevance and requirement of MSI2 function in leukemia was demonstrated by deletion or depletion of MSI2 with a germline gene-trap knockout or shRNAs resulting in reduced leukemogenesis in a CML-BC model^25,26^, whereas forced overexpression of MSI2 and BCR-ABL or NUP98-HOXA13 leads to a more aggressive form of CML^26^ or myelodysplastic syndromes^28^, respectively. *MSI2* is upregulated 10-fold as CML progresses to blast crisis state in patients and shRNA-mediated *MSI2* silencing blocks propagation of both CML-BC and AML cell lines^25,26^. Additionally, *Msi2* was shown to be required for leukemic stem cells (LSC) in a retroviral transplantation MLL-AF9 model of AML^8,30^. We and others have found that MSI mediates its function as an RNA binding protein controling translation of its target RNAs^8,25,30–32^.

Overall, MSI’s requirement in myeloid leukemia makes it an attractive therapeutic target in leukemia and in other malignancies^33^. RNA-binding proteins are often considered “undrugabble” targets due to their lack of well-defined binding pockets for RNA and their absence of enzymatic activity. Structurally, the MSI family of RBPs –comprising the MSI1 and MSI2- contain two highly conserved RNA-recognition motifs (RRMs) in the N-terminal region and a Poly-A Binding Domain (PABP) at the C-terminal region^34^. It is known that RRM1 is the determinant for RNA binding specificity whereas RRM2, mainly adds affinity^35^. The minimal binding consensus described for RRM1 mouse MSI1 is r(GUAG)^36^ and it is known that MSI it also preferentially binds UAG-containing sequences in human and *Drosophila*^35,37^. Here, we describe the identification and characterization of Ro 08–2750. Using biochemical and structural approaches, we find that Ro binds to the MSI2 RRM1 RNA-binding site and inhibits MSI RNA-binding activity and regulation of downstream oncogenic targets. Furthermore, we demonstrate that Ro 08–2750 has efficacy in inhibiting leukemogenesis by *in vitro* and *in vivo* models of myeloid leukemia.

## RESULTS

### Ro 08–2750 (Ro) binds to MSI2 and inhibits its RNA-binding activity

In order to identify a putative MSI inhibitor, we previously performed a fluorescence polarization (FP)-based screen using recombinant MSI1 and MSI2 and a consensus target RNA with a library of 6,208 compounds^38^. We selected Ro 08–2750 (Ro) based on its RNA-binding inhibition of both MSI1 and MSI2^38^. MSI2 RNA-binding inhibition was confirmed by FP (*IC*_50_ of 2.7 ± 0.4 μM) (Fig. 1a) We then used a chemiluminescent Electrophoresis Mobility Shift Assay (EMSA) to quantify MSI2-RNA complexes in vitro. GST-MSI2 bound a MSI RNA, which was competed with the addition of unlabeled RNA and by increasing concentrations of Ro (Fig. 1b and 1c). To confirm the direct interaction of Ro with MSI2 protein, we performed Microscale Thermophoresis (MST) assays with GST-MSI2 and found that the small-molecule interacted with a *K*_D_ of 12.3 ± 0.5 μM (Fig. 1d). RNA-recognition motif 1 (RRM1) of MSI2 also interacted with a similar affinity (Extended Data Fig. 1a) suggesting that the binding was localized to this domain. Similarly, when incubated in the presence of GST-MSI2 and a MSI RNA oligo, Ro could still compete with RNA and bind to MSI2 with a *K*_D_ of 27.5 ± 2.6 μM. We recently found that SYNCRIP, another RNA binding protein, shares MSI2 target RNAs and is also required in leukemia^4^. SYNCRIP has RRMs that are evolutionarily related to MSI’s (with RRM1 and RRM2 sharing 33% and 57% of the residues involved in RNA-binding with MSI2’s RRM1 and 2, respectively) (Extended Data Fig. 1b) Ro showed a 19.2-fold lower *K*_D_ for SYNCRIP than for MSI2 (236.0 ± 167.1 μM, Fig. 1d), indicating selectivity toward MSI2.

**Figure 1.**
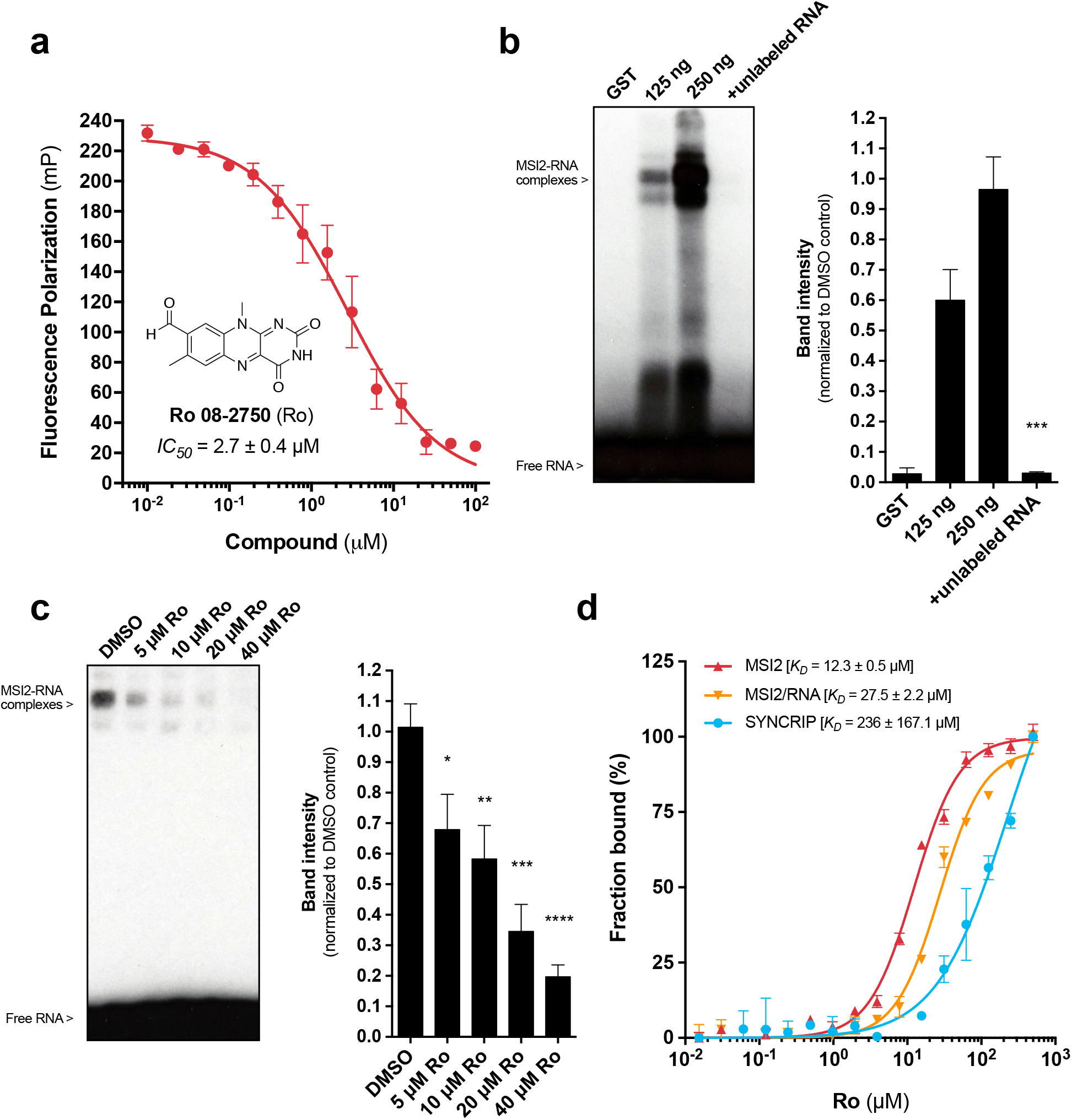
Ro 08–2750 (Ro) is a novel selective MSI RNA-binding activity inhibitor. (a) Fluorescence polarization secondary validation of Ro 08–2750 (Ro) *IC*_50_ for MSI-RNA binding inhibition in 384-well format. Seven independent experiments performed in duplicate ± standard error mean (s.e.m.) are shown; (b) Representative Electrophoresis Mobility Shift Assays (EMSA) for GST- and GST-MSI2 proteins (125 and 250ng) using biotinylated-RNA oligo in the absence or presence of unlabeled RNA (*left*); quantification of MSi2-RNA complexes of at five independent experiments ± s.e.m. is shown in bar graph (*right*); (c) EMSA for GST-MSI2 (125ng) in the presence of increasing concentrations of Ro (5 to 40 μM); quantification of RNA-protein complexes of at least four independent experiments ± s.e.m. is shown in bar graph (right); (d) Microscale Thermophoresis (MST) assay showing interaction of Ro with GST-MSI2, GSTMSI2/RNA complexes or the RRM-RBP control GST-SYNCRIP. Ro concentrations ranged from 0.0153 to 500 μM. Affinity (*K*_D_) values ± s.e.m. (μM) of three independent experiments are shown as percentage of fraction bound. For (b) and (c): two-tailed Paired *t*-test; \**p*<0.05; \*\**p*<0.01, \*\*\**p*<0.005, \*\*\*\**p*<0.001.

### Ro interacts with the RNA recognition site of MSI2 RRM1

To study how Ro interacts with the MSI2 protein, we obtained the crystal structure of *apo* human MSI2 RRM1 at 1.7Å resolution (**Extended Data Table 1**, RCSB PDB accession code 6DBP). This structure allowed us to perform docking analysis to identify a putative binding mode (Fig. 2a, b and Extended Data Fig. 2a). Based on Ro’s ability to compete for MSI-RNA complexes, we hypothesized that the binding site is likely to be shared with the RNA binding site. A closer look at the residues involved in putative Ro binding interactions revealed F66 and R100 as crucial amino acids participating in a stacking interaction with the planar tricyclic structure of the small-molecule (Fig. 2b). Also, the NH backbone group from F97 formed a stabilizing H-bonding with the oxygen from the aldehyde moiety (Fig. 2b, c). A 2D representation of the interacting partners showed R100 forming a π-cation interaction and K22 as a putative amino acid forming an H-bonding with the opposite ring of Ro structure (Fig. 2c). To confirm these putative interactions, we performed site-directed mutagenesis on the full-length MSI2 protein by mutating the F97 or the main three potential residues involved in Ro binding (F66, F97 and R100) to alanine. MST interaction assays showed a nearly 7-fold decrease in affinity (measured *K*_D_ 69.5 ± 14.7 for F97A versus 10.5 ± 0.3 μM for wild-type) for the single mutant. More dramatically, the triple mutant (F66/F97/R100) was incapable of binding Ro, confirming our hypothesis that Ro binds at the RNA-interacting site and can compete for it. We further validated if the mutations of these three residues also disrupt RNA binding by MST: wild-type and F97A possess equivalent RNA binding affinities, whereas the triple mutant F66/F97/R100 only partially ablated RNA binding (*K*_D_ > 50 μM), (Extended Data Fig. 2b).

**Figure 2.**
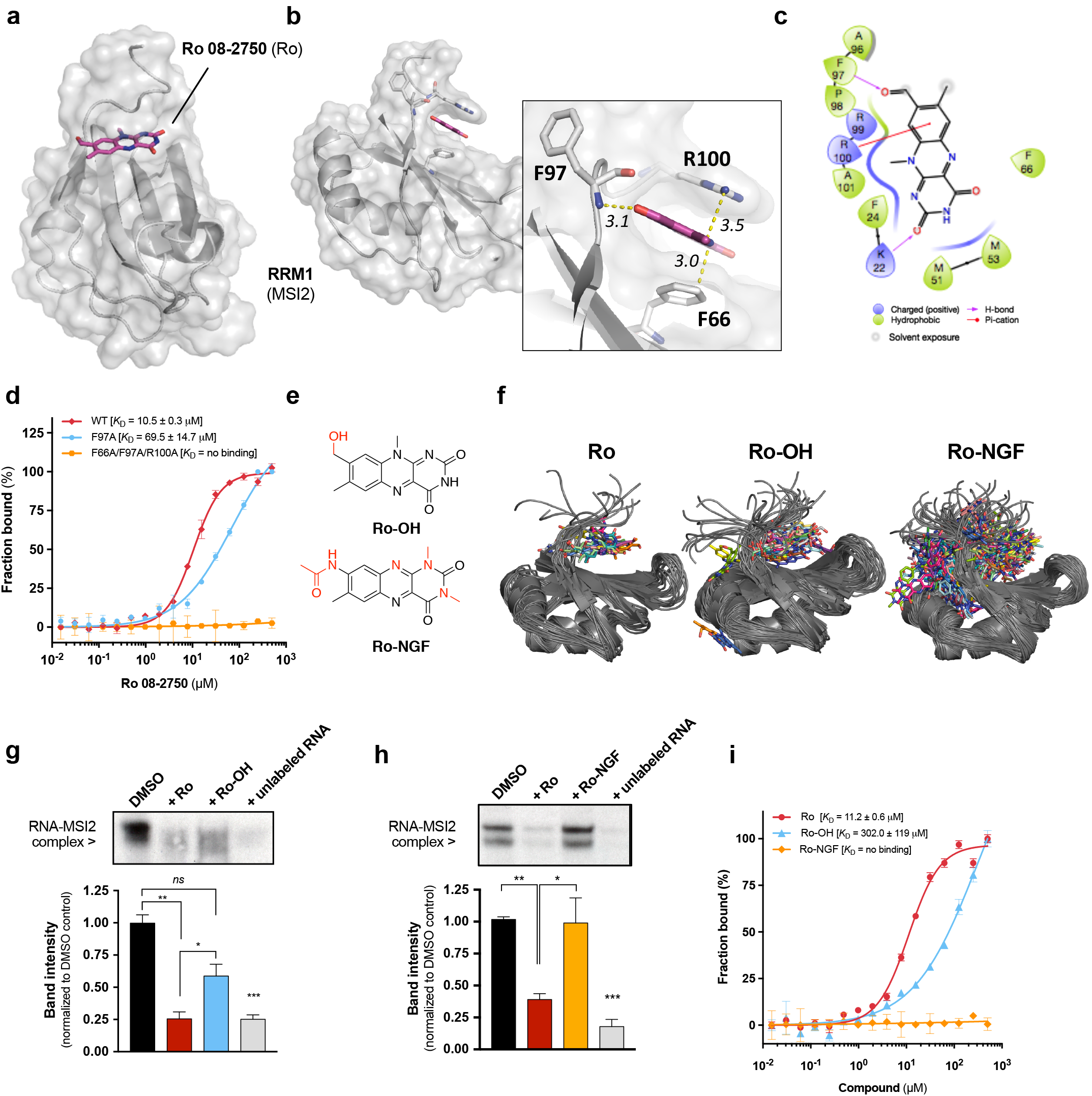
Ro 08–2750 interacts with the RNA-recognition motif 1 (RRM1) of MSI2 and its analogs show minimal or null residual activity. (a) Global front view of the docked Ro 08–2750 (Ro) molecule in the RNA-binding site of human MSI2 RRM1 based on the X-ray diffraction crystal structure obtained at 1.7Å resolution (RCSB PDB 6DBP); (b) Lateral and close up (*inset*) view of Ro showing the most relevant interaction residues (F66, F97 and R100) and the distances (Å) between them and Ro closest atoms; (c) 2D representation of residues involved in Ro binding showing F66 (hydrophobic stacking), K22 (H-bonding), F97 (H-bonding with the backbone) and R100 (π-cation interaction) from RRM1 as main interaction partners; (d) Microscale Thermophoresis (MST) assay showing affinity of interaction of Ro with full-length GST-MSI2 WT (*red*), GST-MSI2 F97A (*cyan*) and GSTF66A/F97A/R100A (*orange*). *K*_D_ values ± standard deviation (μM) of at least three independent experiments are shown as percentage of fraction bound; (e) Chemical structures of Ro analogues used in (f), (g), (h) and (i) panels. Ro-NGF (high affinity Neural Growth Factor –NGF- inhibitor, *K*_D(NGF)_ = 1.7x10^-6^ M) and Ro-OH (reduced form of Ro); (f) The cluster centers for Ro (*left*), RoOH, (*center*) and Ro-NGF (*right*), derived using regular spatial clustering with a ligand RMSD cutoff of 1Å. Ro-NGF (*right*) showing a much larger number of clusters than Ro 08–2750 (*left*) or RoOH (*center*). (g) Representative EMSA for GST-MSI2 (125ng) in the absence (DMSO) or presence of Ro (20 μM), Ro-OH (20 μM) or unlabeled RNA oligo (1 μM) and quantification of RNA-protein complexes of at least three independent experiments (bar graph, *below*); (h) Representative EMSA for GST-MSI2 (125ng) in the absence (DMSO) or presence of Ro (20 μM), Ro-NGF (20 μM) or unlabeled RNA oligo (1 μM) and quantification of RNA-protein complexes of at least three independent experiments (bar graph, *below*) ± s.e.m.; (i) MST assays showing interaction of Ro, Ro-OH and Ro-NGF with GST-MSI2 WT. Drug concentrations ranged from 0.0153 to 500 μM. *K*_D_ values ± standard deviation (μM) of at least three experiments are shown as percentage of fraction bound; For (g) and (h), two-tailed Paired *t*-test; *ns*, not significant, \**p*<0.05, \*\**p*<0.01, \*\*\**p*<0.005.

To further test structure activity relationships, we obtained two Ro related molecules (Ro-NGF and Ro-OH). The first analog, Ro-NGF, was selected to determine if Ro’s activity was related to its anti-NGF activity, as previously described^39^ because this compound showed the highest affinity (*K*_D [NGF]_ = 1.7 μM) for NGF in its compound series (**Extended Data Table 2**). The second analog, Ro-OH, a reduced form of the aldehyde to an alcohol, contained a single alteration to the Ro aldehyde moiety (Fig. 2e and Extended Data Figs. 2c, 3a, b). Alchemical free energy calculations showed computed binding free energies (ΔG_bind_) for the three ligands (Ro, Ro-OH and Ro-NGF) in a similar range, with a slightly higher affinity predicted binding for Ro and Ro-OH (−5.5 and −6.1 *vs* −5.1 kcal/mol for Ro-NGF) (Extended Data Fig. 4a). Both MSI2 protein and ligands adopted a conformationally heterogeneous ensemble of binding poses, with the protein-ligand complex predicted to undergo a slight conformational change for Ro and Ro-OH upon binding (Extended Data Fig. 4b). Free energy calculations for all three small-molecules suggest that Ro-NGF adopts a much more diverse set of conformations (as measured by conformational clustering of the fully-interacting alchemical state) than Ro-OH or Ro (Fig. 2f). Ro showed the fewest clusters, with the top three clusters accounting for 92.7% of the sampled configurations (Extended Data Fig. 4c). Ro-OH showed a larger number of clusters, with the four clusters accounting for 49.1% of sampled configurations, indicating a greater degree of heterogeneity than Ro (Extended Data Fig. 4d). Ro-NGF displayed an even greater degree of heterogeneity, showing a large number of low population of clusters (data not shown). Further structural analysis of our docked model suggests Ro-OH lacking the R100 π-cation interaction and Ro-NGF in a displaced position from the RNA-binding core (Extended Data Fig. 2e, f, g, h) as compared to Ro, despite similar interacting residues. To experimentally validate these predictions, we performed EMSA of GST-MSI2 competing Ro-OH and Ro-NGF with RNA, comparing potency with Ro and unlabeled RNA as positive controls. Accordingly, whereas Ro-OH showed partial (∼30–40%) but significantly poorer inhibition than Ro (65–75%, *p*<0.05), Ro-NGF showed no ability to displace MSI2-RNA complexes (Figure 2g and 2h). These results were further confirmed by FP assay with Ro-OH inhibiting with 12.5-fold less potency than Ro, and Ro-NGF failing to inhibit of RNA-binding activity (**Supplemental Figure 2d**). Furthermore, in MST assays, Ro-OH showed a 27-fold lower affinity than Ro (*K*_D_ 302.0 ± 119 μM for Ro-OH versus 11.2 ± 0.6 μM for Ro) for GST-MSI2, whereas Ro-NGF failed to demonstrate any interaction (Figure 2i). Thus, our structural and biochemical experimental data support the conclusion that Ro and MSI2 interact via the RRM/RNA binding site and that the drug can displace RNA from its binding site, thus likely inhibiting MSI-related translational regulation.

### Ro 08–2750 demonstrates therapeutic efficacy in murine MLL-AF9 leukemic cells

To test the MSI-inhibitory effect of Ro in a murine AML of leukemia, we used MLL-AF9 expressing leukemic BM cells from secondary transplants previously established in the lab^8^. We first assessed the cytotoxicity effects of Ro and the two analogs against these leukemia cells. Consistent with an on-target effect on MSI inhibition and in agreement with the RNA-binding activity inhibition assays, Ro effectively inhibited leukemia cell proliferation (half-effective concentration, *EC*_50_ = 2.6 ± 0.1 μM). By comparison, the analogues that failed to interact with MSI2 had a diminished effect (Ro-OH *EC*_50_ = 21.5 ± 0.8 μM; Ro-NGF > 50 μM), suggesting that the antiproliferative effect is due to the ability of Ro to inhibit MSI2 RNA binding-activity (Fig. 3a). Treatment of cells with Ro resulted in an increase in the myeloid and granulocyte markers (Mac1 and Gr1, respectively) at 5 μM dose and 48h treatment as seen by both flow cytometry (Fig. 3b) and morphologically by Eosin Y and Methylene Blue/ Azure A staining (Fig. 3c). When we assessed apoptosis at different time points, we found a significant increase in the Annexin V + population as early as 8h (both at 5 and 10 μM) with the highest increase at 48h and 10 μM Ro (Fig. 3d and Extended Data Fig. 5).

**Figure 3.**
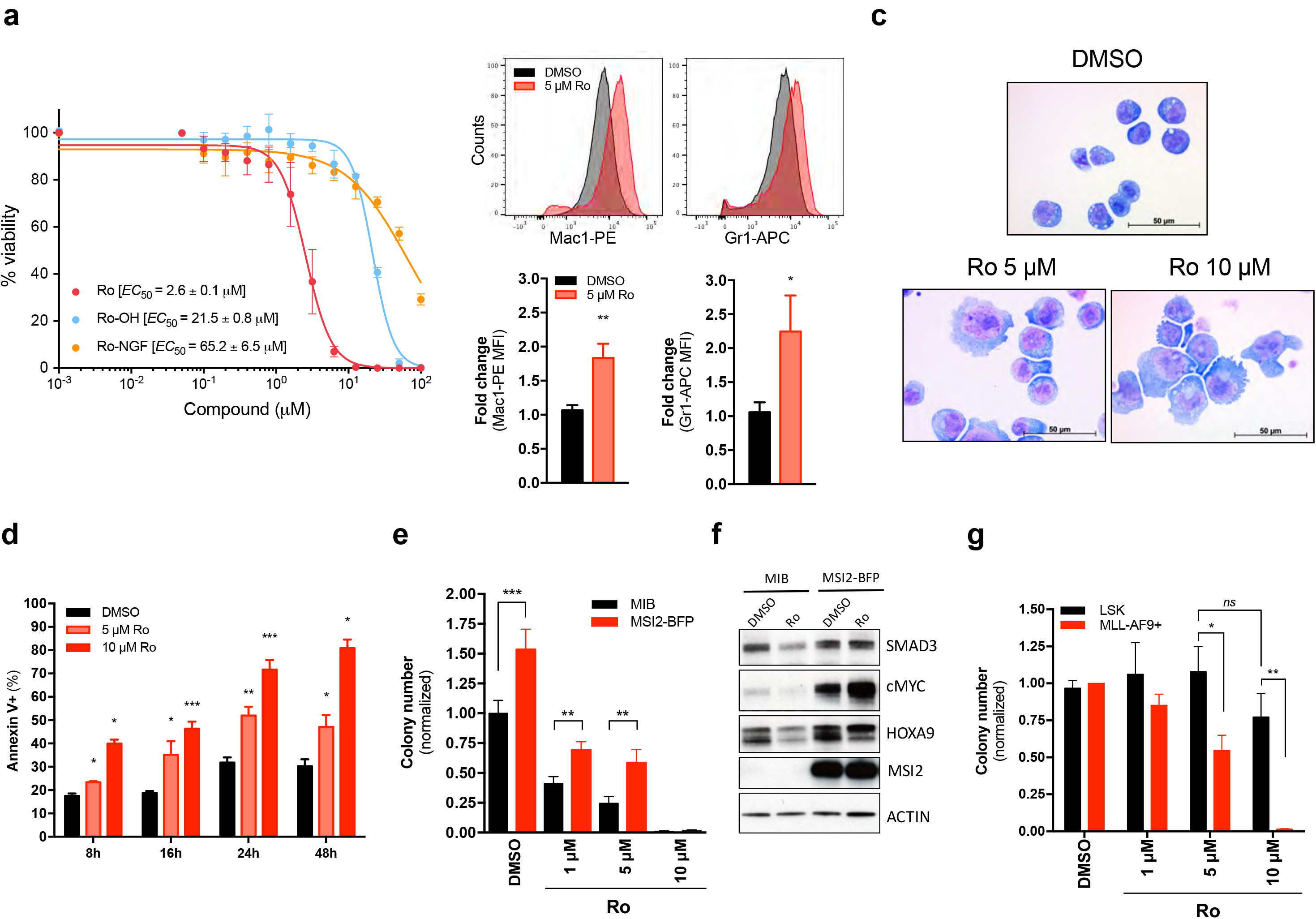
Ro 08–2750 treatment leads to preferential increase in differentiation and apoptosis in murine MLL-AF9 leukemic cells compared to Lin^-^Sca^+^ cKit^+^ (LSK) cells. (a) Cytotoxicity assay (Cell-Titer Glo^®^) of Ro (*red*), Ro-OH (*cyan*) and Ro-NGF (*orange*) in MLLAF9+ BM cells. 50% Effective Concentration (*EC*_50_) values, average of at least three independent experiments ± standard deviation are shown. (b) Flow cytometry representative histograms of DMSO (*grey*) and 5 μM Ro (*red*) treated MLL-AF9+ BM cells showing myeloid differentiation markers (Mac1 and Gr1); bar graphs (*below*) show average (fold change increase) ± standard error mean of three independent experiments, performed in triplicate. Paired *t*-test, \**p*<0.05; \*\**p*<0.01. (c) Representative immunocytochemistry images of cytospun MLL-AF9+ BM cells control (DMSO) or Ro treated (5 and 10 μM) and stained by Eosin Y and Methylene Blue/ Azure A. Scale, 50 μm. (d) Apoptosis analysis by Annexin V+ (% population) for MLL-AF9+ BM cells cultured in absence (DMSO, *black*) or presence of Ro 5 μM (*light red*) or 10 μM (*red*). Results represent at least three independent experiments ± s.e.m.. (e) Colony Formation Unit (CFU) assay of MLL-AF9+ BM cells transduced with MSCV-IRES-BFP (MIB, control) or MSCV-IRES-MSI2-BFP (MSI2-BFP) retroviral vectors. Results represent the average ± s.e.m. of colony numbers of at least five experiments performed in duplicate. (f) Representative immunoblot of MLL-AF9+ BM MIB (*black bars*) and MSI2-BFP (*red bars*) cells (used in panel **e**) after DMSO or 10 μM Ro treatment for 4h. β-ACTIN, loading control. (g) CFU assay of Lin^-^Sca^+^cKit^+^ (LSK) versus MLL-AF9+ BM cells demonstrates Ro 08–2750 therapeutic window. Results represent the average ± s.e.m. of colony numbers of three experiments performed in duplicate. Two tailed Paired *t*-test (b, d, e and g), \**p*<0.05, \*\**p*<0.01, \*\*\**p*<0.005.

We then assessed how MSI2 overexpression affected the plating capacity of MLL-AF9 BM cells in culture in the absence or presence of Ro. MSI2 overexpressing cells formed 50% more colonies than control cells transduced with an empty vector (MIB). Treatment of cells with Ro resulted in reduced colony formation in control cells by > 50% and ∼75% at 1 μM and 5 μM concentrations, respectively. MSI2-overexpressing leukemia cells however showed increased resistance to these doses (Fig. 3e). Of note, we assessed MSI2 translational targets^8,31^ in these cells by immunoblotting and we found that Ro treatment reduced protein abundance of SMAD3, c-MYC and HOXA9 in control cells, whereas the levels of these proteins remained unaffected in cells that overexpressed MSI2 (Fig. 3f). Indicating a potential therapeutic window between normal and malignant cells, Ro abolished MLL-AF9+ BM colony formation at concentrations that did not affect the plating efficiency of normal Lin-Sca+cKit+ (LSK) cells (Fig. 3g),.

### Ro 08–2750 treatment inhibits survival of human AML cell lines and patient cells

To determine if Ro also has activity against human myeloid leukemia, we first tested cytotoxicity effects of the small-molecule and the two analogs in MOLM13 (AML, MLL-AF9+) and K562 (CML-BC, BCR-ABL+) cell lines, both known to require MSI2 function^4,26^. Consistent with our previous data in MLL-AF9 cells, we observed that in both these leukemia cell lines, Ro demonstrated anti-proliferative effect (*EC*_50_ ∼8 μM), whereas the two analogs (Ro-OH and Ro- NGF) revealed a > 4.5-fold weaker potency. Ro induced myeloid differentiation and apoptosis in both K562 and MOLM13 cells based on flow cytometry and by morphology (Fig. 4b–d and **Extended Data Fig. 6a-c** and **6b**). Plating activity was > 80% inhibited at the 20 μM Ro dose in the human AML cell lines (Fig. 4e). Additionally, Ro demonstrated differential sensitivity in three AML patient samples (**Extended Data Table 3**) colony plating assays compared to normal human CD34+ cord blood cell (> 50% inhibition in colony numbers at 5 μM comared to only a modest reduction at 20 μM Ro, Fig. 4f). These results indicate that Ro can induce differentiation and apoptosis in primary human AML cells and spare normal stem cells up to 2 x *EC*_50_ Ro.

**Figure 4.**
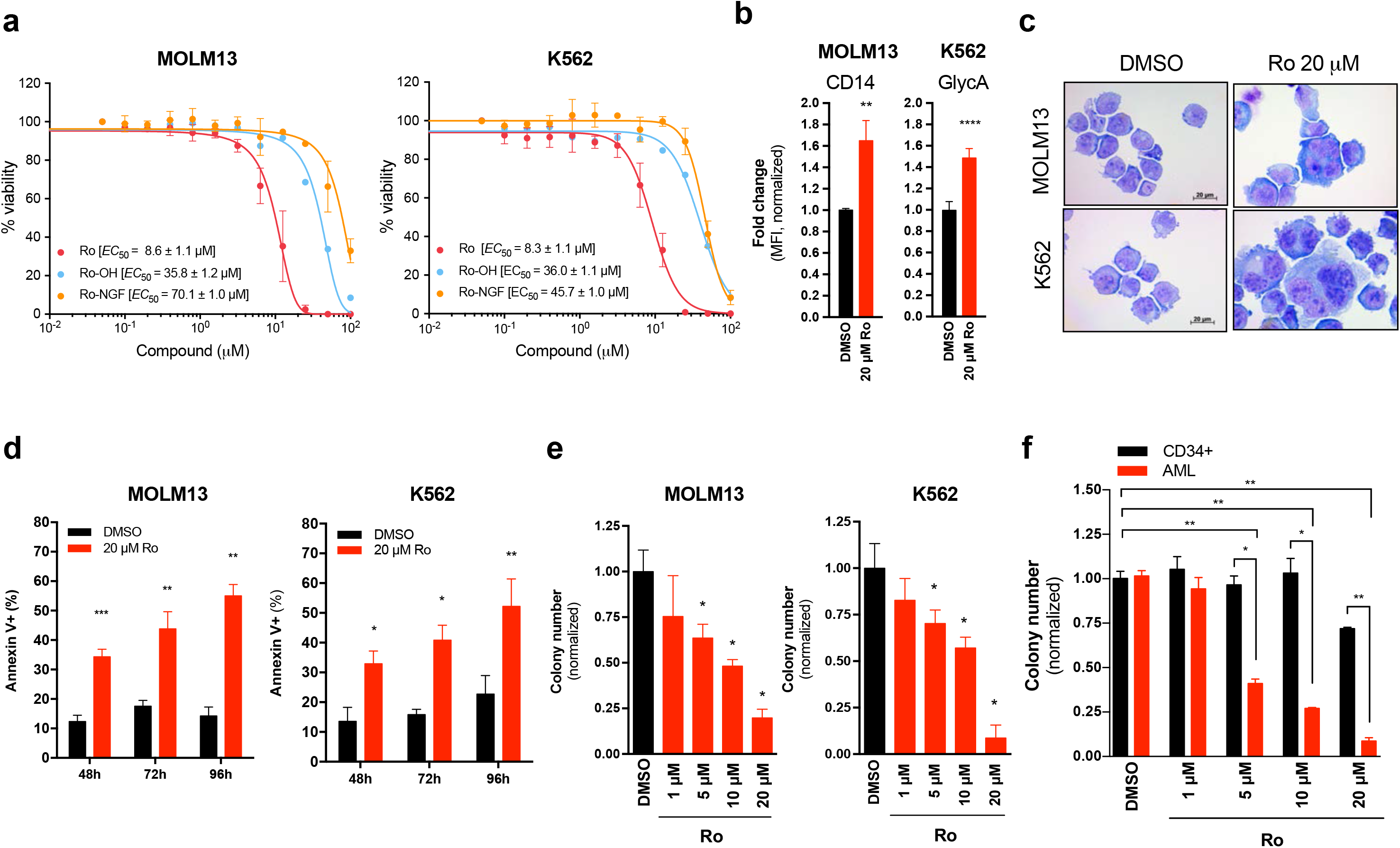
Ro 08–2750 treatment inhibits survival of human AML cell lines and patient cells. (a) Cytotoxicity assay (Cell-Titer Glo^®^) of Ro, Ro-OH and Ro-NGF in MOLM13 and K562 cells. *EC*_50_ values average of three independent experiments ± standard deviation is shown. (b) Mean Fluorescence Intensity (MFI) fold changes of CD14 (myeloid marker, MOLM13) and CD235a (Glycophorin-A; erythroid marker, K562) after 48h treatment with DMSO (control, *black bars*) or Ro 20 μM (*red bars*). Data is normalized to DMSO control cells. Representative histograms are shown in *Extended Data Figure 6a*. (c) Representative immunocytochemistry images of cytospun MOLM13 and K562 cells treated for 48h with DMSO (control) or Ro 20 μM and stained with Eosin Y and Methylene Blue/ Azure A. Scale, 20 μm. (d) Apoptosis analysis by Annexin V+ (% population). MOLM13 and K562 were cultured in DMSO (*black bars*) or in the presence of Ro 20 μM (*red bars*) for the indicated times and Annexin V positivity and 7AAD was measured. Results represent three independent experiments ± standard deviation. (e) CFU assay of MOLM13 and K562 in the presence of Ro 08–2750 at different concentrations (1, 5, 10 and 20 μM). Data is shown as average colony numbers (normalized to DMSO control) ± s.e.m. of at least three independent experiments. (f) CFU assay of cord-blood derived CD34+ HSPCs and AML patient BM cells. Data is shown as average colony numbers (normalized to DMSO) ± s.e.m. of three different blood donors for CD34+ and three independent AML patients. Two tailed Paired *t*-test (DMSO vs Ro treated, unless indicated with lines); \**p*<0.05; \*\**p*<0.01; \*\*\**p*<0.005, \*\*\*\**p*<0.001.

### Ro 08–2750 inhibits binding of MSI2 to its RNA targets and exhibits gene signature from MSI2 depleted cells

To further investigate the effect and mechanism of action of Ro, we initially performed RNA immunoprecipitation (RNA-IP with FLAG) experiments on K562-MIG (empty vector) and K562-FLAG-MSI2 (MSI2 overexpressing) cells (Fig. 5a). After incubating the drug at 10 μM (*∼EC*_50_) for 1 hour with the cells, we could detect a significant decrease in MSI2 mRNA binding targets (*TGFBR1, cMYC, SMAD3, CDKN1A*) (Fig. 5b). These data suggest that Ro can block MSI2 binding to target mRNAs in a cellular context at a short time-point.

To globally assess the proximal effect of Ro treatment on the transcriptional program, we then performed RNA-sequencing on MOLM13 and K562 cells after 4 hours of treatment. Ro incubation resulted in modest but significant gene expression changes in both the MOLM13 and K562 AML cells (59 upregulated, 221 downregulated and 111 upregulated, 164 downregulated, respectively; FDR<0.05), (**Extended Data Tables 4–5**). Most importantly, this Ro signature enriched for the gene expression profiling after shRNA mediated depletion of MSI2 in CML-BC (AR-230 and LAMA84) and AML cell lines (THP1 and NOMO-1) (Fig. 5c)^26^. To annotate the functional pathway overlap with Ro treatment in both cell lines and MSI2 shRNA depletion, we performed gene-set enrichment analysis (GSEA)^40^ on all 4,733 curated gene sets in the Molecular Signatures Database (MSigDB, http://www.broadinstitute.org/msigdb) combined with 92 additional relevant gene sets from our experimentally derived or published hematopoietic self-renewal and differentiation signatures^31,40^. Interestingly, we observed an overlap of MSI-associated signatures from our previous dataset and an enrichment with MSI1 direct mRNA targets from the intestine (**Extended Data Tables 7–12** and Extended Data Fig. 7a)^4^. Moreover, we observed a ∼70% overlap of the functional pathways between each individual cell line and the pathways altered after shRNA depletion of MSI2 (Fig. 5d). Among these shared pathways, 76% (543 out of 717) overlapped in MOLM13 compared to K562 cells treated with Ro, which included c-MYC, mRNA related and leukemia associated gene sets (Fig. 5d and **Extended Data Table 12**). Thus, Ro treatment after a short administration recapitulated a large portion of the MSI2-associated gene expression program.

To determine how Ro affects previously determined MSI targets, we treated both K562 and MOLM13 cells with increasing concentrations of Ro (up to 20 μM at 4 hours). In previous studies, MSI was demonstrated to maintain the protein levels of TGFβR1, c-MYC, SMAD3 and HOXA9^8,31^ while suppressing P21 abundance^41,42^. Consistent with this, we observed a significant and dose dependent reduction of TGFβR1, c-MYC, SMAD3, HOXA9 and an increase in the protein abundance of P21, while the non-target control β-ACTIN remained unchanged (**Fig. 5d** and **5e**). Additionally, Ro could inhibit MSI2 targets in a time-dependent manner with c-MYC, a short half-life protein, being reduced in 1 hour of treatment (**Fig. 5f** and **5g**). In support of Ro altering translation of specific MSI2 targets but not generally inhibiting global translation, we found equivalent global protein synthesis after drug treatment as assessed by *O*-propargyl-puromycin incorporation (Extended Data Fig. 7b). As previously noted by RNA-sequencing, there were modest effects on the mRNA expression of MSI2 targets by qPCR (Extended Data Fig. 7c) suggesting that Ro mainly influences its direct targets through a post-transcriptional mechanism. Thus, these results support our hypothesis that Ro acts in the MSI-related translational program.

**Figure 5.**
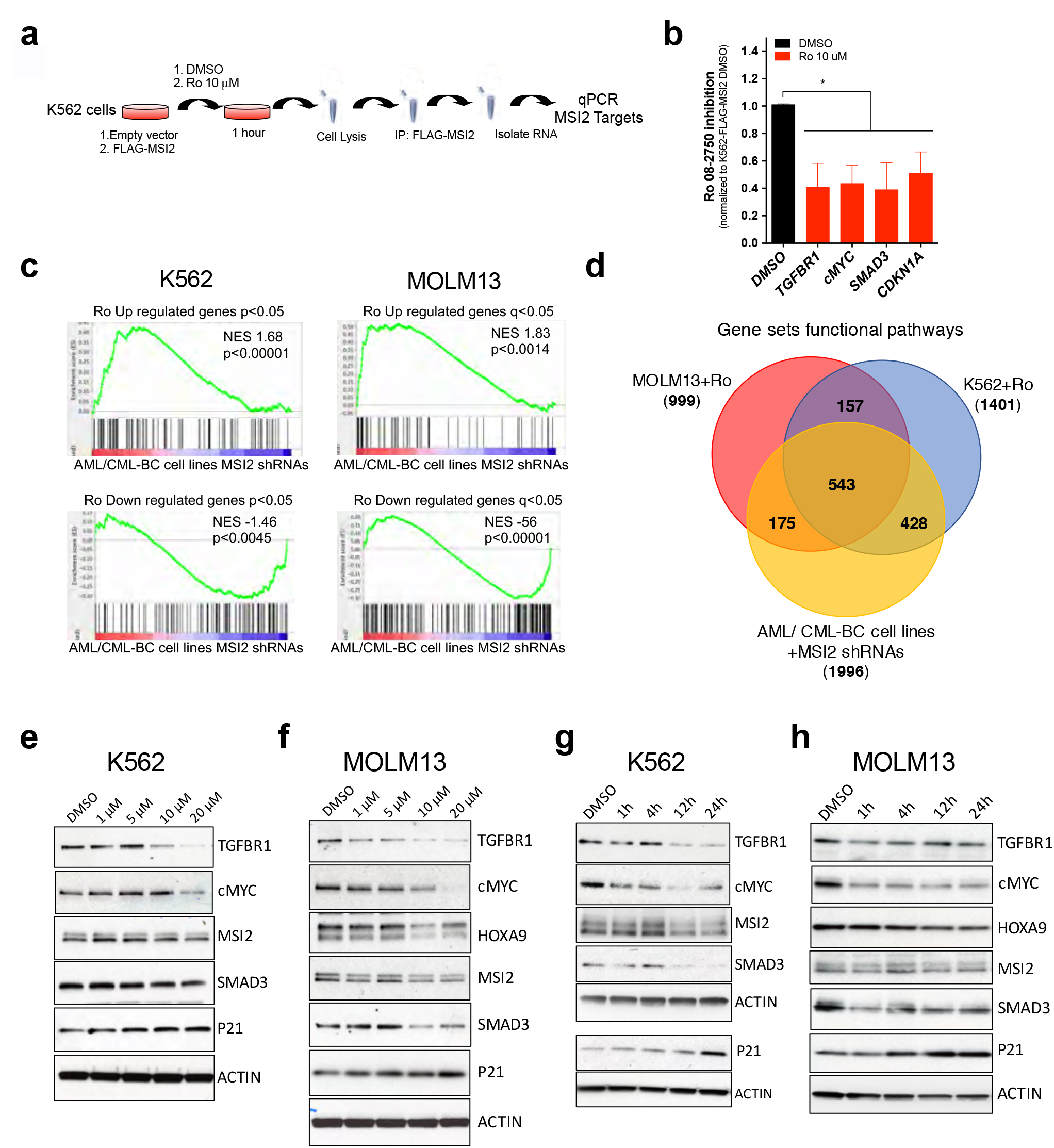
Ro 08–2750 treatment resembles gene signature from MSI2 depleted cells and demonstrates inhibition of MSI2 target translation. (a) Scheme of RNA-immunoprecipitation (IP) protocol followed with K562-MIG (MSCV-IRES-GFP) or FLAG-MSI2 overexpresing cells. (b) Ro 08–2750 inhibitory effect in the RNA-IP enrichment of MSI2 mRNA targets in K562-FLAG-MSI2 versus K562-MIG after 1h treatment at 10 μM. Data is shown as average of inhibition effect (normalized to DMSO cells) ± s.e.m. of four independent experiment. (c) Up-regulated and down-regulated gene sets obtained by RNA-seq analysis after 20 μM Ro 4h treatment in K562 and MOLM13 cells showing identical signature as previously obtained using shRNA against MSI2 in CML-BC and AML lines^26^. (d) Venn diagram showing gene Set Enrichment Analysis (GSEA) overlap between MOLM13 (*red*), K562 (*blue*) (after 20 μM Ro 4h treatment) and AML/CML-BC cell lines MSI2 depleted with shRNAs (*yellow*) from^26^. Bold values inside brackets below each grup are total gene sets numbers. (e) Representative immunoblot for K562 treated with Ro at different concentrations (1, 5, 10 and 20 μM) for 4h showing expression of MSI2 targets. HOXA9 is not expressed in this BCR-ABL+ (CML-BC) leukemia cell line. (f) Representative immunoblot for MOLM13 treated with Ro at different concentrations for 4h showing expression of MSI2 targets. (g) Representative immunoblot for K562 treated with Ro 20 μM at different time points (1, 4, 12 and 24h) showing expression of same MSI2 targets as in panel (e). P21 and β-ACTIN from a different representative gel are shown. (h) Representative immunoblot for MOLM13 treated with Ro 20 μM at different time points showing effect on MSI2 targets.

### Ro 08–2750 inhibits leukemogenesis in an in vivo MLL-AF9 model of myeloid leukemia

Finally, we sought to determine if Ro has activity in vivo using an aggressive murine MLL-AF9 murine leukemia model. Acute treatment of Ro (4h and 12hr) reduced c-KIT protein abundance and intracellular c-MYC (Fig. 6a-c). To determine if Ro treament could affect disease burden we next treated a second cohort of animals and monitored them for disease progression for 19 days after transplantation (Fig. 6d). Ro administration every 3 days was well tolerated (Extended Data Fig. 7a, b, c) demonstrating little to no weight loss and equivalent red blood cells and platelets counts compared to control group. When control mice succumbed to disease (day 19 post-transplantation), we assessed the disease in both groups and found a significant reduction in spleen weights (Fig. 6e), white blood cell counts (Fig. 6f) and c-MYC levels compared to the controls (Figure 6g). These data provide the feasibility that targeting MSI in vivo could have therapeutic efficacy in AML.

**Figure 6.**
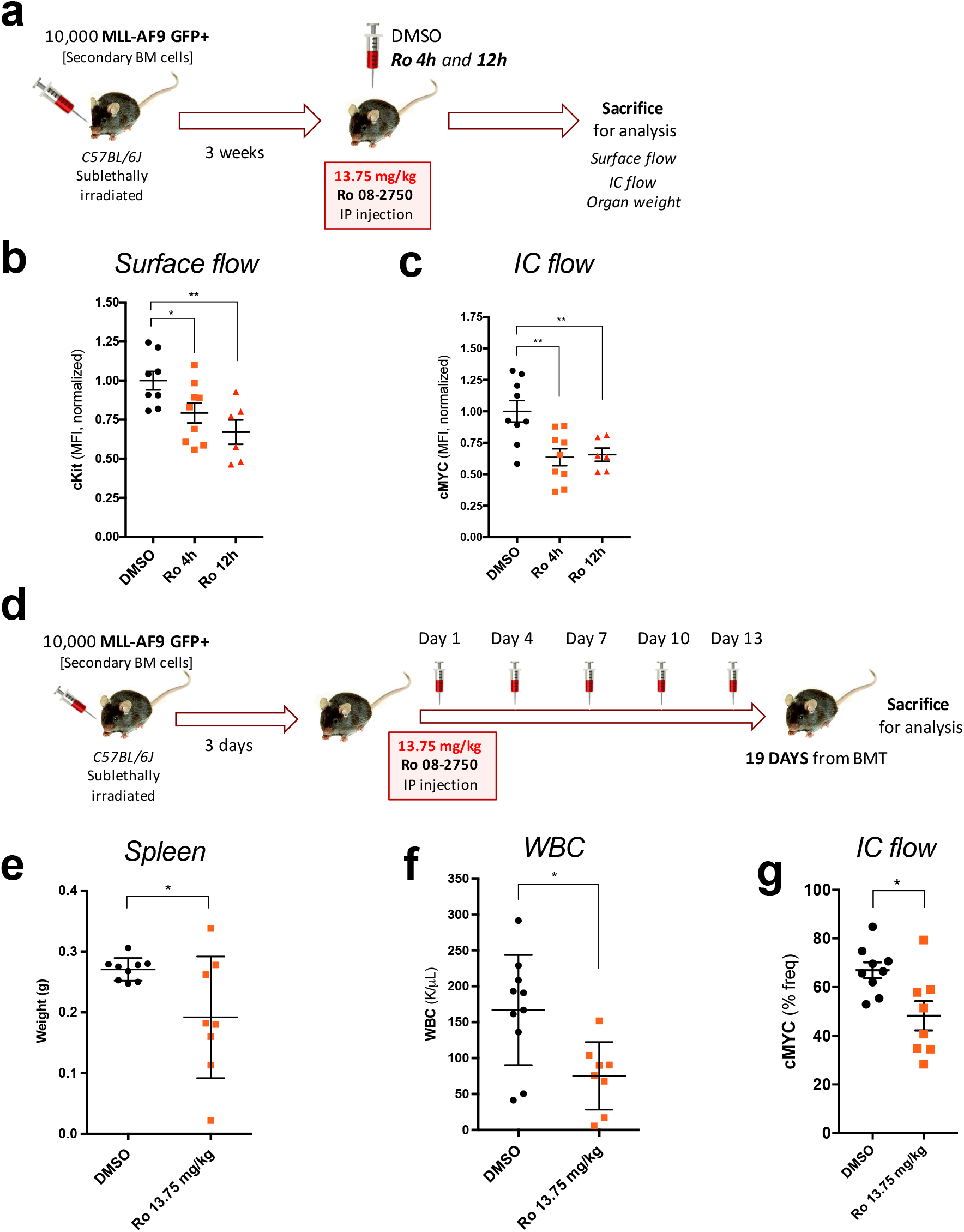
Ro 08–2750 demonstrates efficacy in inhibiting leukemogenesis in short-time and long-term treatment in a MLL-AF9 *in vivo* model. (a) Scheme of pharmacodynamics marker experiments with Ro short-time points performed with MLL-AF9+ secondary BM cells. 10,000 MLL-AF9 GFP+ cells were transplanted and, after 3 weeks, mice were injected with DMSO or Ro (13.75 mg/kg) and were sacrificed for analysis after 4h and 12h (b) Surface flow analysis of c-Kit receptor in spleen cells of Ro at 4h and 12h *versus* DMSO treated mice. Results are represented as MFI of cKit-PE-Cy7 normalized to DMSO group. Each data point is an independent treated mouse. Mean ± s.e.m. is shown. (c) Intracellular (IC) flow analysis of c-MYC expression in spleen cells of Ro at 4h and 12h *versus* DMSO treated mice. Results are represented as MFI of c-MYC normalized to DMSO group. Each data point is an independent treated mouse. Mean ± s.e.m. is shown; (a-c, DMSO and Ro 4h, *n* = 9; Ro 12h, *n* = 6). (d) Scheme of *in vivo* Ro treatment in MLL-AF9+ model of myeloid leukemia. 10,000 MLLAF9 GFP+ cells were transplanted and after 3 days, mice were injected with DMSO or Ro 13.75 mg/kg (in DMSO) intraperitoneally (IP) at days 1, 4, 7, 10 and 13 (one day *on*, two days *off* drug). At day 19 of treatment, mice were sacrificed for organ weight and flow cytometry analysis of disease burden and MSI2 target, c-MYC. (e) Spleen weights at time of sacrifice. Results are represented in weight (g) and each data point represents an individual DMSO or Ro treated mouse. (f) White blood cell (WBC) counts (K/μL) at time of sacrifice. Each data point represents an individually treated mouse. (g) Intracellular (IC) flow analysis of c-MYC expression in spleen cells of Ro vs DMSO treated mice. Results are represented as% frequency (% freq) of c-MYC+ cells. Each data point is an independent treated mouse. Mean ± s.e.m. is shown. (d-g, DMSO, *n*=9; Ro, *n*=8). For all graphs, Unpaired *t*-test; \**p*<0.05, \*\**p*<0.005.

## DISCUSSION

Inhibiting MSI RNA-binding activity could represent a novel therapeutic avenue in both hematological malignancies and solid cancers. Our previous FP-based screen identified compounds that inhibit MSI binding to RNA^38^. Here, we characterize Ro 08–2750 as a first selective MSI inhibitor with biochemical, structural and cellular validation linking the compound to the inhibition of the MSI program. Ro falls in the low micromolar range of activity, in line with other RBP associated inhibitors^43–47^. We validated Ro as a MSI2 RNA-binding inhibitor with biophysical and biochemical assays by utilizing a high-resolution crystal structure of the MSI2 RRM1. Our newly developed computational molecular modeling algorithm and docking analysis, allowed to predict and validate the key MSI2 residues that were critical for the interactions with Ro in the RNA binding site. Both our novel crystal structure and the computational tools will be useful for the discovery and development of small-molecule RBPs inhibitors. We found that a single chemical reduction of Ro decreased its activity in both in biochemical and in vitro cell based assays. Utilizing a related compound with high affinity binding to NGF, we found that it no longer bound MSI2 and poorly inhibited leukemia cell growth. Further studies involving medicinal chemistry with heterocycle isoalloxazines or pteridine-derived compounds could help identifying more selective and potent MSI-inhibitors.

Other groups have identified agents that have putative MSI1 inhibitory activity. A natural phenol extracted from cottonseed ((-)-gossypol) was shown to reduce MSI1 to bind RNA^45^ but this interaction was not validated by structure-activity relationships. Of note, (-)-gossypol has been considered to be a pan-active compound that has hit in multiple HTS screens^48–50^ and assigned to have activity against Bcl-2^51^. MSI1 activity was also inhibited by γ-9 monounsaturated fatty acids (e.g. oleic acid), allosterically binding and inducing a conformational change that prevents RNA to bind^52^. It remains unclear if (-)-gossypol or oleic acid have a more broad RNA binding protein inhibitor activity as they were not directly tested against any other RBPs^43,45,52^. We found that Ro could demonstrate differential binding activity to MSI2 compared to SYNCRIP, Ro’s effect on colony formation and direct targets could be rescued by MSI2 overexpression. Moreover, we observed a strong enrichment for the MSI2 shRNAs gene expression signature, associated functional pathways, inhibition of MSI2 binding of target mRNAs and reduced abundance of MSI2 direct targets after Ro treatment. In contast to other general translational inhibitors Ro did not alter global translation^53,54^. These data suggest that Ro could be used to probe the acute effects of MSI inhibition in a variety of cellular contexts and cancer models.

It is also important to note that Ro inhibits both MSI1 and MSI2 and although MSI1 is expressed at low levels in myeloid leukemia it could still be blocking residual MSI1 activity. Moreover, in other models such as the intestine where both factors act redundantly^13^, dual inhibition could provide a powerful therapeutic strategy. Of note based on the close conservation of the RRMs between the two proteins it might be challenging to design MSI1 or MSI2 selective inhibitors.

We demonstrated a therapeutic index for Ro in human AML patient samples versus cord-blood derived CD34+ human stem and progenitor cells. Despite the challenges for in vivo administration, we reduced the disease burden in an aggressive MLL-AF9 leukemia model and decreased MYC levels without overt toxicity. Interestingly, it has previously been shown that MSI2 can contribute to chemotherapeutic resistance in different cancer models^42,55,56^. Future studies could examine if combination therapies could provide additional clinical benefit.

This study identifies and characterizes Ro 08–2750 as the first compound selectively inhibiting the oncogenic RNA-binding activity of MSI in myeloid leukemia. It will be important to use this compound (or other chemical derivatives) to test their efficacy in other cancer models and on MSI function related to normal physiology. We suggest that Ro provides the rationale for developing more potent compounds with improved clinical utility for the treatment of cancers that are dependent on the MSI family. Additionally, as there are hundrends of RRM containing RNA binding proteins, Ro targeting an RRM motif to block RNA activity represents a valuable proof of concept for the general inhibition of these class of RNA regulators. Thus, we provide a framework to identify and test novel RNA binding protein inhibitors in cancer.

### Methods

#### Purification and culture of cord blood derived HSPC-CD34+ cells

Mononuclear cells were isolated from cord blood using Hetarstach solution (6% Hetastarch in 0.9% NaCl) and Ficoll-Hypaque Plus density centrifugation. CD34+ Hematopoietic Stem and Progenitor Cells (HSPCs) were subsequently purified by positive selection using the Auto MACS Pro Separator and isolation kit (Miltenyi) and were cultured in Iscove’s modified Dulbecco’s medium (IMDM, Cellgro), 20% BIT 9500 medium (Stem Cell Technologies) supplemented with SCF (100 ng/ml), FLT-3 ligand (10 ng/ml), IL-6 (20 ng/ml) and TPO (100 ng/ml) as the basic culture. All cytokines were purchased from Peprotech, NJ.

#### Isolation and viral transduction of murine MLL-AF9 leukemia and normal cells

Tibia, femurs, pelvis, and arm bones from leukemia or C57BL/6 wild type mice (10–12 weeks old) were harvested, crushed, filtered, and subjected to red blood cell lysis (Qiagen). To isolate c-Kit^+^ cells, bone marrow cells were incubated with anti-CD117 microbeads (Miltenyi Biotec), according to manufacturer’s instructions, and then subjected to positive selection using autoMACS Pro Separator. For MLL-AF9 BM cells, previously thawed vials from secondary transplants (Park et al. 2015) were used. All murine cells were cultured and transduced in RPMI with 10% FBS and cytokines SCF (10 ng/ml), IL-3 (10 ng/ml), and IL-6 (10 ng/ml) and GM-CSF (10 ng/ml). For MSI2 overexpression, cells were spinfected with viral supernatant containing MSCV-IRES-BFP or MSI2-IRES-BFP contructs (see Cloning section).

#### Colony forming unit (CFU) assays

10,000 leukemic MLL-AF9 BM cells or c-Kit enriched normal stem cells (Lin-Sca-Kit+) were plated on methylcellulose-based culture media (methocult) GFM3434 (Stem Cell Technologies). Colonies were scored every five days for leukemia cells and every seven to nine days for normal c-kit-enriched bone marrow cells. For human cells, 5,000 of the leukemia cel lines K562 (CMLBC) or MOLM13 (AML) and 10,000 of HSPCs CD34+ or AML patient cells were plated (in duplicate) in methylcellulose (MethoCult^TM^ H4434 Classic, Stem Cell Technologies). CFU colonies in HSPCs CD34+ were scored 14 days after seeding. AML patient cells characteristics are shown in **Extended Data Table 3**.

#### Flow cytometry

To monitor the differentiation status, 200K MLL-AF9 BM cells DMSO or Ro treated (during 8, 16, 24, 48h) were stained with the following antibodies: anti-CD11b (Mac1)-PE (clone M1/70, #101208, BioLegend), anti-Ly-6G (Gr1)-APC (clone RB6–8C5, #17–5931-82, eBioscience), and anti-CD117 (c-Kit)-APC-Cy7 (clone 2B8, #105826, BioLegend). For the human cell lines differentiation, we used two panels: (1) anti-CD14-PE (clone M5E2, #555398, BD Pharmingen), anti-CD13-APC (clone TUK1, #MHCD1305, Life Technologies); (2) anti-CD71-APC (clone CY1G4, #334104, BioLegend), anti-CD235a (Glycophorin A)-PE (clone YTH89.1, #MA5–17700, Invitrogen). All samples were stained for 20min in the dark, washed once with PBS 1X and re-suspended in RPMI+ 2% FBS for analysis. For intracellular flow cytometry detection of cMYC, 1–2x10^6^ cells were fixed in 2% paraformaldehide for 15 minutes, washed 2 times with 1X PBS and permeabilized with cold methanol and kept at –80 until use. For the staining, cells were washed twice in 1X PBS and stained in 100 μl final volume. c-MYC (5605, Cell Signaling Technology non-labelled primary antibody was incubated at 1/200 dilution for 1h and labelled donkey anti-rabbit Alexa Fluor 568 (#A10042, Invitrogen) or goat anti-rabbit Alexa Fluor 647 (#A21245, Invitrogen) were used at 1/400 for 20–30 minutes. Cells were washed once with PBS 1X and re-suspended in RPMI+ 2% FBS for analysis. All flow cytometry analysis was performed in a LSRII or LSR Fortessa (BD Biosciences) and data was graphed by using FlowJo^TM^ version 10.4.

#### Morphological analysis

After the appropriate time of Ro treatment (or DMSO in controls) in culture, 1.5x10^5^ MLL-AF9 BM or human leukemia cells (K562 and MOLM13) were washed once with 1X PBS, counted and centrifuged onto slides for 5 minutes at 500 rpm and air-dried for 24h prior to Richard-Allan Scientific Three-Step Stain Staining Set (Thermo Scientific) based on Eosin Y and Methylene Blue/ Azure A and mounted with Permount solution (Fisher). Cell morphology was evaluated by light microscopy at 400X magnification (Zeiss Imager M-2, equipped with AxioCam ERc 5s).

#### Apoptosis measurements

Apoptosis measurements were taken by MUSE^TM^ Cell Analyzer (Millipore) using the MUSE^TM^ Annexin V and Dead Cell Assay Kit (Millipore) as recommended by the instructions from the manufacturer. Dot plots showing viability versus Annexin V+ cells are shown in ***Extended Data Figures 4*** and ***5***.

#### *In vivo* transplantation of leukemia cells and Ro 08–2750 administration

10,000 of MLL-AF9 BM secondary mouse leukemia cells previously obtained^1^ were injected retro-orbitally into female C57BL/6 (10–12 weeks old) recipient mice that had been sublethally irradiated at 475 cGy. Drug administration (Ro 08–2750, 13.75 mg/Kg, DMSO) was performed by intraperitoneal injections (50 μL, top tolerated DMSO volume) 3 weeks after BM transplants (when showing signs of disease) for pharmacodynamic experiments (see Fig. 6a), and 3 days after BM transplant for in vivo long-term studies (see Figure 6d). Mice weight were monitored every day to check for toxicity. All animal studies were performed on animal protocols approved by the Institutional Animal Care and Use Committee (IACUC) at Memorial Sloan Kettering Cancer Center.

#### Fluorescence Polarization (FP) to assess RNA-binding activity inhibition

To validate RNA-binding activity inhibition by Ro 08–2750 and derivatives (Ro-OH, Ro-NGF) we used Fluorescence Polarization (FP) based assay in as previously described in 384-well format for dose-response curve studies^2^. As previously, the RNA oligo used (Cy3-C_9_-[spacer]-rGUAGUAGU, Integrated IDT Technologies) contained 2 MSI motifs (G**UAG**U) and was 8-nucleotides long, optimal to minimize background and unspecific interactions. Differently, here manual pipetting was used to plate the reagents and the FP reading was performed in a BioTek Synergy Neon Plate Reader (High-Throughput Screening Resource Center, Rockefeller University).

#### Binding affinity quantifications by MicroScale Thermophoresis (MST)

For binding affinity studies of RNA and small-molecules to proteins of interest, purified recombinant GST-MSI2 WT, F97A and F66A/F97A/R100A mutants and GST-SYNCRIP were NT647-labeled using an amine-coupling kit (NanoTemper Technologies). Runs were performed at a concentration range of 50–125 nM (MSI2 and mutants) and 60 nM (SYNCRIP) to get optimal fluorescence signal using an LED power of 40–50% in a red laser equipped Monolith NT.115 (NanoTemper Technologies) (High-Throughput Screening Resource Center, Rockefeller University). Prior to each run, protein preparations were diluted in MST buffer (50 mM HEPES, 100 mM NaCl, 0.05% Tween-20, pH 7.4) and aggregation was minimized by centrifuging the solutions at 15,000 rpm for 10 minutes. GST-proteins or GST-protein/RNA complexes (15 min pre-incubation) were mixed with increasing concentrations of small-molecules (0.015 to 500 μM) or RNA (0.0015 to 50 μM) and loaded onto 16 Premium Coated capillaries. The RNA oligo used (rGUAGUAGUAGUAGUA, Integrated IDT Technologies) contained 4 MSI motifs (G**UAG**U) and was 15-nucleotides long. The MST measurements were taken at RT and a fixed IR-laser power of 40% for 20 seconds per capillary. GraphPad Prism was used to fit the normalized data and determine apparent *K*_D_ values, represented as percent of fraction bound.

#### Chemiluminescent Electrophoresis Mobility Shift Assays (EMSA)

An EMSA approach to assess MSI2-RNA complexes and the inhibitory effect of small-molecules was set up by using LightShift Chemiluminescent RNA EMSA kit (Thermo Scientific). In brief, GST-MSI2 (125–250 ng) was preincubated with DMSO or the small-molecule (typically 20 μM final concentration) during 1h at RT in EMSA buffer 1X RNA EMSA binding buffer (10 mM HEPES, 20 mM KCl, 1 mM MgCl_2_, 1 mM DTT, Thermo Scientific) supplemented with 5% glycerol, 100 μg/mL tRNA and additional 10 mM KCl. After this period, 40 nM of biotinylated-RNA (biotinrGUAGUAGUAGUAGUA, Integrated IDT Technologies –same as for MST-) was added to the mixture (20 μL final volume) and incubated another 1h at RT. During this second incubation period, a 4–20% TBE polyacrylamide gel (BioRad) was pre-run at 100V for 30–45min in cold 0.5X TBE (RNAse free). 5 μL of 5X loading buffer was added to the 20 μL reaction and loaded into the pre-run TBE gel and voltage set at 100V. Samples were electrophoresed until 3/4 of the length of the gel. Samples were then transferred in 0.5X TBE at 350–400 mA for 40 min. Membranes were then crosslinked with UV-light crosslinking instrument (UV Stratagene 1800) using Auto-Cross Link function. Membranes were either stored dry for development next day or developedusing the detection biotin-labeled RNA chemiluminescence kit (as indicated by the manufacturer) (Thermo Fisher) and Hyperfilm ECL (GE Healthcare).

#### Cloning, expression, and purification of GST tagged proteins

Human full-length MSI2 was cloned into the retroviral backbone pMSCV-IRES-BFP (MIB) vector (a gift from Dario Vignali; Addgene plasmid # 52115) by Custom DNA Constructs (University Heights, Ohio) introducing a 5’Flag tag and using BamHI and EcoRI restriction sites. Human full-length MSI2 was previously cloned into pGEX6P3 as described^2^. RNA-recognition motif 1 (RRM1) from human MSI2 (nucleotides #64–270, NM_138962.2) was subcloned into empty pGEX6P3 using EcoRI and NotI restriction sites. Human SYNCRIP (hnRNP-Q variant 3, NM_001159674.1) was subcloned into empty pGEX6P3 (GE Healthcare) by introducing a 5’Flag sequence (5’-ATGGATTACAAGGATGACGACGATAAG-3’) and using SalI and NotI sites. GST-Flag-MSI2 wild-type (WT), Flag-MSI2 mutants (F97A, F66A/F97A/R100A), GST-RRM1 and GST-Flag-SYNCRIP recombinant proteins were produced in BL21 (DE3) competent cells (Agilent Technologies, Santa Clara, CA) as previously reported for MSI2 WT^2^. Here, GST-SYNCRIP protein needed higher content of NaCl (250 mM) in the 1X PBS dialysis step and final buffer for optimal storage and performance in the biochemical and biophysical assays performed.

#### Site-directed mutagenesis

To perform site-directed mutagenesis into pGEX6P3-Flag-MSI2 construct and express the corresponding recombinant GST-MSI2 mutants, we used QuikChange Lightning and Multi Site-Directed Mutagenesis Kit (#210513 and #210518, Agilent Technologies). The primers were designed using QuickChange Primer Design (https://www.genomics.agilent.com/primerDesignProgram.jsp) and were the following: F66A (Fwd: 5’-GCTCCAGAGGCTTCGGTGCCGTCACGTTCGCAG-3’, Rev: 5’-CTGCGAACGTGACGGCACCGAAGCCTCTGGAGC-3’; F97A/ R100A (Fwd: 5’-AGACGATTGACCCCAAAGTTGCAGCTCCTCGTGCAGCGCAACCCAA-3’, Rev: 5’-TTGGGTTGCGCTGCACGAGGAGCTGCAACTTTGGGGTCAATCGTCT-3’) and R100A (using F97A mutant construct as template) (Fwd: 5’-CCAAAGTTGCAGCTCCTCGTGCAGCGCAACCCA-3’, Rev: 5’- TGGGTTGCGCTGCACGAGGAGCTGCAACTTTGG-3’). PCR reactions and cloning were perfomed as indicated by the manufacturer (Agilent Technologies).

#### Human MSI2 RRM1 recombinant protein production

GST-RRM1 protein was initially produced in BL21 (DE3) competent cells (Agilent Technologies, Santa Clara, CA) as previously reported for MSI2 WT^2^. Here, the cell lysate of 4L initial culture was centrifuged at 15,000 rpm for 1h and the resulting volume applied to a XK16/20 column pre-packed with Glutathione Sepharose 4 Fast Flow connected to an AKTA Prime FPLC (GE Healthcare). To obtain the RRM1 optimal prep for the crystal preparation, the collected fractions containing GST-RRM1 (in 50 mM Tris-HCl, 20 mM reduced L-Glutathione) were pooled and dialyzed against PreScission Protease Buffer (50 mM Tris-HCl, 150 mM NaCl, 1 mM EDTA, 1mM DTT, pH 7.5). GST tag was then cleaved with PreScission Protease overnight at 4°C. Pure RRM1 fractions were obtained through size exclusion chromatography (HiLoad Superdex 75, GE Healthcare) and concentrated with a 3K Amicon Ultra Centricon (Millipore).

#### Crystallization and structure determination

A final concentrated MSI2 RRM1 pure protein preparation (> 98% by coomassie) at 2 mg/mL in 50 mM Tris-HCl, 150 mM NaCl, 1 mM EDTA, 1mM DTT, pH 7.5 was crystallized by sitting drop vapor diffusion. A 1 uL of protein solution was mixed with an equal volume of precipitant solution containing 100 mM Tris, 200 mM Li2SO4, 25% PEG (pH 8.5). Crystals appeared after two weeks. They were cryoprotected by mother liquor containing 25% glycerol and flash frozen in liquid nitrogen. X-ray diffraction data were collected from single crystals at the Advanced Photon Source beamline 24ID-C at 100 K. Indexing and merging of the diffraction data were performed in HKL2000^3^. The phases were obtained by molecular replacement by PHENIX^4^ using PDB entry 1UAW as the search model. Interactive model building was performed using O^5^. Refinement was accomplished with PHENIX. Data collection and refinement statistics are summarized in **Extended Data Table 1**. The crystal structure has been deposited in RCSB PDB under the accession code 6DBP.

#### RNA purification and quantitative real-time PCR

Total RNA was isolated from 1–2x10^6^ cells dry pellets kept at –80C for less than a week using Qiagen RNeasy Plus Mini kit. cDNA was generated from RNA using iScript cDNA Synthesis (#1708891, BioRad) with random hexamers according to the manufacturer’s instructions. Real-time PCR reactions were performed using a Vii7 sequence detection system. β-*ACTIN* was commonly used to normalize for cDNA loading. Relative quantification of the genes was performed using Power SYBR Mix (2X) and specific primers for *c-MYC, TGFβR1, SMAD3, HOXA9* and *CDKN1A* and the 2^−ΔΔ*C*t^ method as described by the manufacturer.

#### Immunoblot analysis

For immunoblot analysis, Ro treated and DMSO control MOLM13 or K562 cells (routinely at 0.5x10^6^ cells/ mL) were counted and washed twice with cold PBS before collection. 1–5x10^6^ cells were resuspended and lysed in 250 μl of 1X RIPA Buffer supplemented with Protease Inhibitor Tablets (Sigma-Aldrich) buffer for 30min on ice. After centrifugation at 14,000rpm on a top-bench centrifuge, lysate (supernatant) was collected and total protein quantified by BCA (Thermo Scientific). Cell lysates were separated by 4–15% SDS–PAGE and transferred to 0.45 μm nitrocellulose membrane. Membranes were blocked and were blotted overnight (4C) for TGβR1 (ab31013, Abcam, 1:750 dilution), SMAD3 (9523S, Cell Signaling Technology, 1:750 dilution), HOXA9 (07–178, Millipore, for drug dose-dependent experiments and ab140631, Abcam; 1:1,000 dilution for time-course experiments), c-MYC (5605, Cell Signaling Technology; 1:1,000 dilution), P21 (2947S, Cell Signaling Technology, 1:750 dilution), MSI2 (ab76148, Abcam; 1:2,000 dilution) and β-ACTIN-HRP conjugated (A3854, Sigma-Aldrich; 1:20,000 dilution) and developed by Hyperfilm ECL (GE Healthcare) with ECL and pico-ECL reagents (Thermo Scientific).

#### Luminescence-based cytotoxicity assays (*EC*_50_)

10,000 cells (MLL-AF9 BM from secondary transplants or human leukemic cell lines –K562 or MOLM13-) were platted into U-bottom 96-well plates in the presence of increasing concentration of small-molecules (Ro, Ro-OH or Ro-NGF) up to 100 μM (in 1:2 serial dilutions). Cells were cultured for 72h at 37C in a 5% CO_2_ incubator. To read cell viability, Cell-Titer Glo^TM^ kit (Promega) was used. After cooling down cells to RT for 20–30min, 100 μL of the cultured cells were transferred to opaque-white bottom 96-well plates and mixed with 100 μL of Cell-Titer Glo^TM^ Reagent (previously prepared by mixing buffer and powdered substrate). The mixture was incubated for 15min at RT and read using a Synergy H1 Hybrid reader (BioTek) for luminescence. Data was normalized as percentage viability and graphed by non-linear regression curves in Graph Pad PRISM 7.0. K562 and MOLM13 lines were purchased from ATCC, authenticated by Genetica, and tested negative for mycoplasma contamination.

#### RNA immunoprecipitation (RNA-IP)

To assess mRNA enrichment and blocking of protein-binding to mRNA by the small-molecules we performed RNA immunoprecipitation (RNA IP) experiments using Magna RIP RNA-binding protein immunoprecipitation kit (#03–115, Millipore). 25 × 10^6^ K562-MIG or MSI2 overexpressing cells 1h treated with DMSO (control) or Ro μM were used. First, cells were washed with cold PBS and lysed. Five micrograms of mouse anti-Flag (clone M2, #F1804, Sigma-Aldrich) antibody incubated with magnetic beads were used to immunoprecipitate Flag-MSI2 K562 cells. After washing the immunoprecipitated, they were treated with proteinase K. RNA extraction was performed by the phenol–chloroform method, and 200–500 ng of purified RNA was converted to cDNA using the Verso cDNA kit (Thermo Scientific). qPCR was used to validate target mRNAs bound by MSI2 and control cells.

#### *O*-Propargyl-Puromycin incorporation by flow cytometry

Cells were plated at a density of 200,000 cells/ml and pre-treated with DMSO or Ro up to 4h. Then, 50 μM *O*-propargyl-puromycin (OP-Puro; NU-931–05, Jena Bioscience) was added. Control cells were co-incubated with DMSO or Ro and treated with 150 μg/ml cycloheximide for 15 min. Non-OP-Puro treated cells were also used as negative controls for flow cytometry. Cells were washed twice before collection and subjected to processing using the Click-iT Flow Cytometry Assay kit (C10418, Invitrogen) following the manufacturer’s instructions. Labeled cells were analyzed using a BD LSR Fortessa instrument and graphed as Alexa Fluor 647 (AF647) Mean Fluorescence Intensity (normalized to DMSO control treated with OP-Puro).

#### RNA sequencing

Total RNA was isolated from 1x10^6^ dry pellets of K562 and MOLM13 4h treated with DMSO (control) or Ro 20 μM (*n* = 4 for each group) using Qiagen RNeasy Plus Mini kit and the quality assessed on a TapeStation 2200 (Agilent technologies). QuantSeq 3′ mRNA-Seq Library Prep Kit FWD (Lexogen, Vienna Austria), supplemented with a common set of external RNA controls, according to manufacturer’s recommendations (ERCC RNA Spike-In mix, ThermoFisher Scientific, #4456740). An in-house pipeline was used for read mapping and alignment, transcript construction and quantification of data generated by sequencing (HiSeq 2000, NYGC, NY, USA). This procedure was done in the Epigenetics Core from MSKCC. RNA-seq data has been deposited to GSE114320 and can be viewed for reviewers only: https://www.ncbi.nlm.nih.gov/geo/query/acc.cgi?acc = GSE114320.

#### Synthesis of Ro-OH by reduction of Ro 08–2750 aldehyde

To a cooled (0 °C) slurry of Ro 08–2750 (19 mg, 0.070 mmol) in anhydrous MeOH (1.9 mL) was added LiBH_4_ (32 mg, 1.5 mmol) in portions over 5 min. The slurry turned from bright orange to dark brown, then dark green within 10 min. The reaction mixture was removed from the ice bath and allowed to warm to rt (22 °C) over 2 h. Reaction progress was monitored by LC-MS (5–95% MeCN in H_2_O). Four portions of LiBH_4_ (10 mg, 0.04 mmol) were added every 12 h until the reaction was complete. The reaction was quenched with AcOH (10 mL) and filtered. The solids were washed with water (5 mL), MeOH (5 mL), and Et_2_O (5 mL). The solid was collected and dried under vacuum to provide a pale orange solid (7 mg, 26%). Purification by HPLC (5–95% MeCN in H_2_O) afforded the product as an orange solid (3 mg, 16%). The synthesis was adapted from Salach *et al.^6^*

**^1^H-NMR** (600 MHz, DMSO) ´ 11.34 (s, 1H), 7.91 (s, 1H), 7.87 (s, 1H), 5.69 (t, *J* = 4.5, 1H), 4.74 (d, *J* = 4.4, 2 H), 3.99 (s, 3H), 2.38 (s, 3H).**^13^C-NMR** (150 MHz, DMSO) 159.8 (C), 155.4 (C), 150.5 (C), 149.7 (C), 137.4 (C), 133.57 (C), 133.56 (C), 131.5 (CH), 131.0 (C), 112.3 (CH), 60.8 (CH_2_), 31.7 (CH_3_), 17.2 (CH_3_); IR (ATR): 2361, 2341, 1717. **ESI-MS** *m/z* (rel int): (pos) 273.1 ([M+H]^+^, 100).

#### Statistical analysis

Student’s *t* test was used for significance testing in the bar graphs, except where stated otherwise. A two-sample equal-variance model assuming normal distribution was used. The investigators were not blinded to the sample groups for all experiments. *P* values less than 0.05 were considered to be significant. Graphs and error bars reflect means+ standard error of the mean except stated otherwise. All statistical analyses were carried out using GraphPad Prism 7.0 and the R statistical environment.

#### Modeling and System preparation for Computational Modeling

System preparation, modeling, and initial docking calculations were performed using the Schrödinger Suite molecular modeling package (version 2015–4), using default parameters unless otherwise noted. The MSI2 RRM1 protein structure (PDB ID: 6DBP) was prepared using the Protein Preparation Wizard^7^. In this step, force field atom types and bond orders were assigned, missing atoms were added, tautomer/ionization states were assigned, water orientations were sampled, and ionizable residues (Asn, Gln, and His residues) have their tautomers adjusted to optimize the hydrogen bond network. A constrained energy minimization was then performed. All crystallographiclly resolved water molecules were retained.

Potential binding sites were explored and characterized using the SiteMap^8,9^ tool. Ligands with experimental activity and known inactives were docked into putative binding sites using Glide SP^10,11^ to evaluate enrichment of known actives. Best docking scores were for the ‘Ro’ series for the ‘(-)-gossypol’ binding site described by Lan *et al*.^12^ compared to other putative pockets.

Since the receptor may not be in an optimal conformation to bind small molecule inhibitors, induced fit docking^13^ of ligand Ro 08–2750 was performed to this binding pocket. Induced fit docking results were validated with the metadynamics protocol described by Clark *et al*.^14^ In these metadynamics simulations a biasing potential is applied to the ligand RMSD as collective variable. The resulting potential energy surface is evaluated towards how easy a ligand can move away from the initial binding mode. The underlying assumption is that a ligand pose which is closer to the real one has a higher energetic barrier to leave the pose than an incorrect pose. The pose ranked second using the induced fit docking score retrieved the best score from the metadynamics ranking protocol compared to the other induced fit docking poses. This receptor configuration was furthermore tested towards its suitability for a virtual screening by a Glide SP docking of known actives into this pocket. The docking scores using this receptor conformations were better (down to –6.2) compared to the initial protein conformation in the crystal structure. Furthermore, a WaterMap^15,16^ calculation was done for this receptor.

#### Induced Fit Docking of Ro-NGF and Ro-OH compounds

Induced Fit Docking (IFD) was performed against the receptor pose from the selected Ro 08–2750 pose, using Schödinger molecular modeling suite (version 2017–4). Poses for Ro-NGF and Ro-OH, the top and second scored poses respectively, were selected to most closely match the Ro 08–2750 pose.

#### Alchemical Free Energy Calculations

Absolute alchemical free energy calculations were carried out to validate the putative binding poses in a fully-flexible explicitly solvated system. The YANK GPU-accelerated free energy calculation code with the Amber family of forcefields was used for this purpose. Details follow: *System preparation and modeling*. The top poses generated by induced fit docking, as described above, were selected as input protein and ligand poses. Because proteins and ligands were already prepared, they were simply run through the pdbfixer 1.4 command line tool with add-atoms and add-residues set to None to convert residue and atom names to be compatible with Amber tleap.

*Parameterization*. tleap (from the minimal conda-installable AmberTools 16 suite ambermini 16.16.0) was used to solvate the complex in a cubic box with a 12Å buffer of TIP3P water molecules around the protein^17^. The system was parameterized using AMBER’s forcefield ff14sb^18^ and GAFF 1.8^19^. Missing ligand parameters were determined using antechamber^20^. The ligand was assigned charges using the AM1-BCC^21,22^ implementation in OpenEye (OEtoolkit 2017.6.1^23^ through openmoltools 0.8.1).

*Minimization*. Minimization was performed using the implementation of the L-BFGS^24^ algorithm in OpenMM 7.1.1^25^ with a tolerance of 1kJ/mol/nm.

*Production Simulation*. Production simulation was run using YANK 0.19.4^26^ using OpenMMTools 0.13.4. In order to keep the ligand from diffusing away from the protein while in a weakly coupled state, it was confined to the binding site using a Harmonic restraint with an automatically-determined force constant (K = 0.33 kcal/mol/Å^2^). The restraint was centered on the following receptor residues using all-atom selection: 2, 4, 46, 76, 78, and 80. The ligand atoms were automatically determined. The calculation was performed using particle mesh Ewald (PME)^27^ electrostatics with default YANK settings with a real-space cutoff of 9Å. A long-range isotropic dispersion correction was applied to correct for truncation of the Lennard-Jones potential at 9Å. The system was automatically solvated with TIP3P^28^ solvent and four neutralizing Cl^-^ ions, paramterized using the Joung and Cheaham paramters^29^. Production alchemical Hamiltonian exchange free energy calculations were carried out at 300 K and 1 atm using a Langevin integrator (VRORV splitting)^30^ with a 2 fs timestep, 5.0 ps^-1^ collision rate, and a molecular-scaling Monte Carlo barostat. Ro 08–2750 and Ro-NGF were run for 10000 iterations (50 ns/replica) with 2500 timesteps (5 ps) per iteration, while Ro-OH was run for 15000 iterations (75 ns/replica) with 2500 timesteps (5 ps) per iteration. Complex configurations were stored for each replica once per iteration. Replica exchange steps were performed each iteration to mix replicas using the Gibbs sampling scheme described previously^31^. The alchemical pathway was automatically determined for each compound using the YANK autoprotocol protocol trailblazing feature.

*Absolute binding free energy estimates*. Absolute free energies (ΔG) of binding for each compound was estimated using MBAR^32^. Samples were reweighted to a cutoff of 16Å to correct the isotropic dispersion correction to a nonisotropic long-range dispersion. This correction is important to account for the heterogeneous density of protein. To remove the harmonic restraint bias, samples were reweighted to substitute a squared well restraint of radius 10Å.

*Clustering analysis*. The fully interacting trajectory from YANK was extracted to a PDB file, discarding the following number of initial iterations, which came prior to equilibration^33^: 1500 for Ro 08–2750, 1600 for Ro-OH, and 1600 for Ro-NGF. These trajectories were aligned in MDTraj^34^ using only protein backbone atoms. The small molecules were then sliced out and clustered on Cartesian coordinates using the MSMBuilder^35^ implementation of RegularSpatial clustering using a 1Å RMSD cutoff. For the most populated clusters for Ro 08–2750 and Ro-OH, cluster centers were selected and shown with 10 randomly sampled cluster members. Ro-NGF produced a large number of lowly populated clusters with highly heterogeneous binding poses, and were therefore not shown.

*Conformational heterogeneity analysis*. To investigate the conformational heterogeneity in the presence or absence of the ligand, the fully interacting thermodynamic state (corresponding to the holo protein bound to the ligand) and fully non-interacting state (corresponding to the apo protein free of ligand interactions) for all three ligands were extracted using a 4-frame skip, discarding the initial frames as above.

*Code availability*. All Schrödinger project files, YANK simulation inputs, and analysis scripts have been made publicly available (https://github.com/choderalab/musashi).

## Extended Data Figure Legends

**Extended Data Figure 1.**
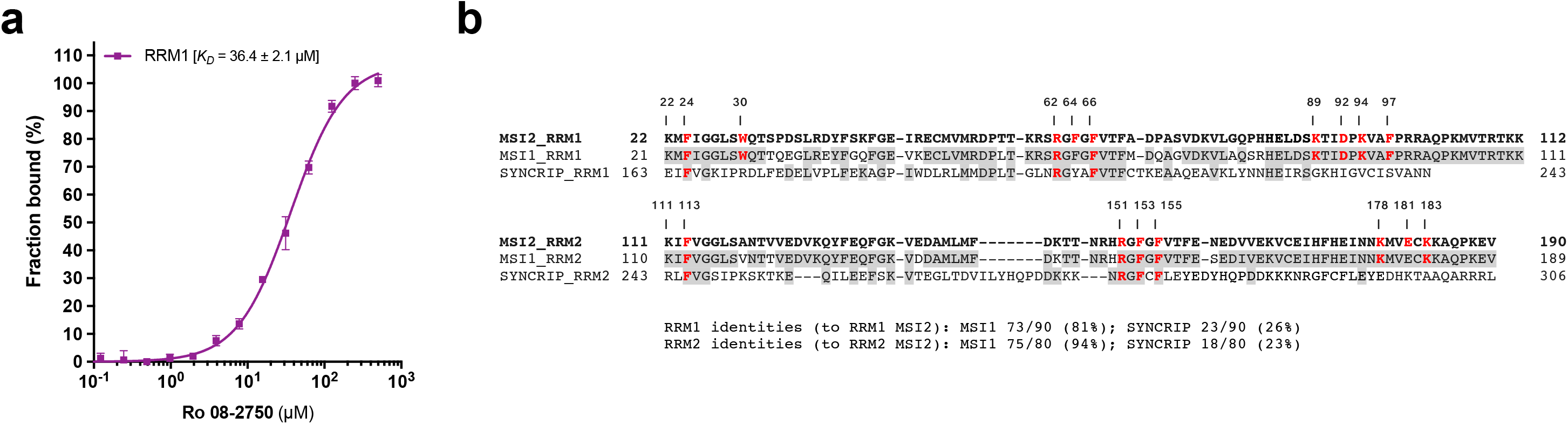
Ro 08–2750 (Ro) binds to RRM1 and SYNCRIP RRM identities. (a) MicroScale Thermophoresis (MST) assay showing interaction of Ro with GST-RRM1 (hMSI2). Ro concentrations ranged from 0.0153 to 500 μM. *K*_D_ values ± s.e.m. (μM) of at least three experiments are shown as percentage of fraction bound. (b) Sequence alignment of RRM1 (*above*) and RRM2 (*below*) of human MSI2, MSI1 and SYNCRIP. Numbers indicate crucial RNA-binding conserved residues (in bold red) in hMSI2 (e.g. F24, corresponding to F23 in hMSI1, F165 in SYNCRIP). Grey highlights indicate conserved amino acids.

**Extended Data Figure 2.**
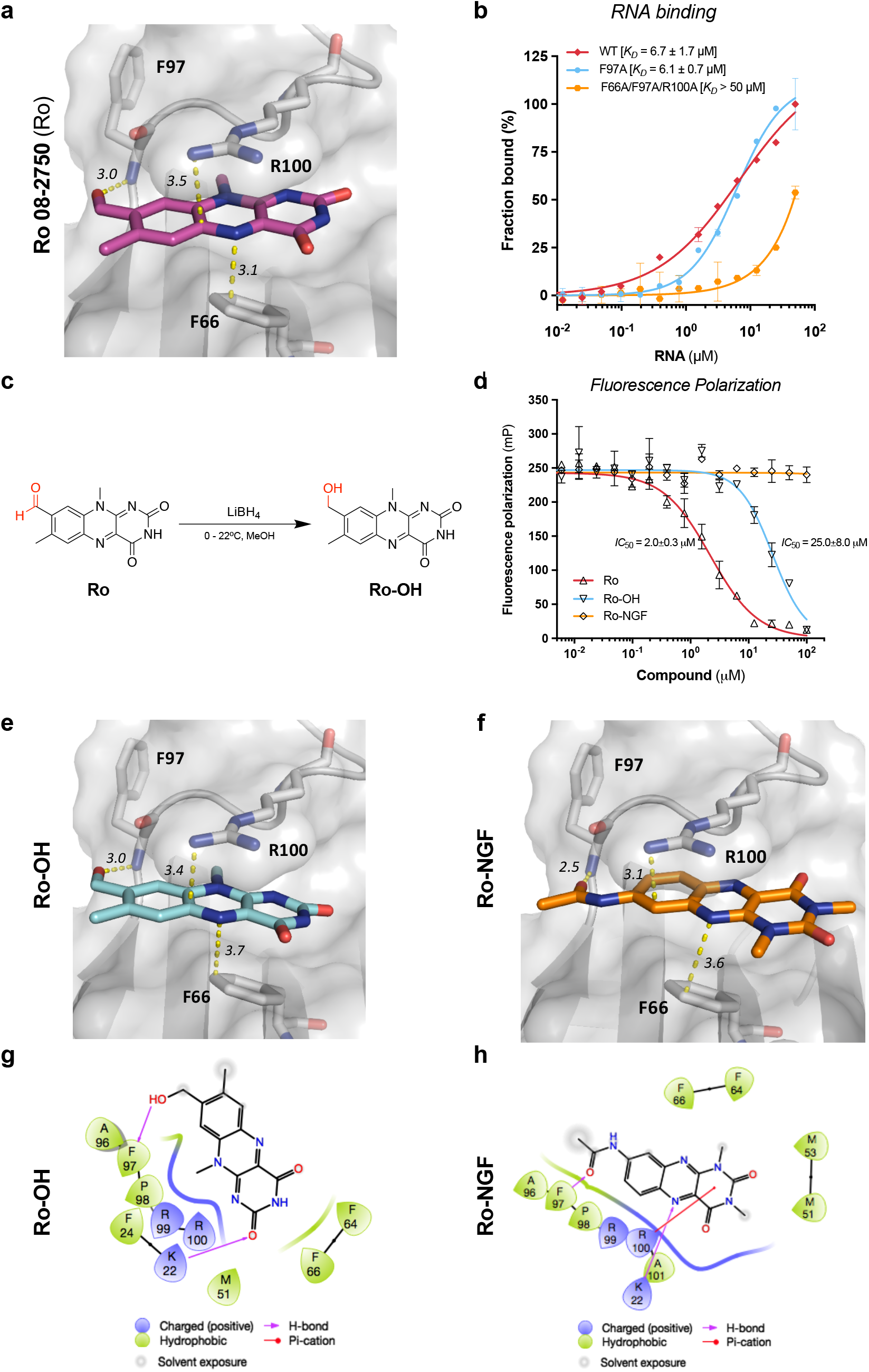
Ro 08–2750 docking and interacting residues in comparison with Ro-OH and Ro-NGF in the RNA-binding site of RRM1. (a) Ro docked in the RNA-binding site of MSI2 RRM1 with interacting residues. Distances shown in Å; (b) MST experiments showing GST-MSI2 WT (*red*), F97A (*cyan*) and Triple (F66A/F97A/R100, *orange*) mutants interaction to MSI2 RNA oligo (4 MSI motifs; 15-nt). *K*_D_ values ± s.e.m. of at least three experiments are shown (μM); (c) Chemical synthesis scheme of Ro-OH from Ro 08–2750 compound (see *Methods*); (d) FP confirmation of Ro, Ro-OH and Ro-NGF MSI2-RNA binding inhibition in 384-well format. *IC*_50_ values of two independent experiments performed in triplicate with s.e.m., 2.0±0.3 μM (Ro, *red*) and 25.0±8.0 μM (RoOH, *cyan*). Ro-NGF (*orange*) showed null inhibition of RNA-binding activity; (e) Docked pose of Ro-OH in the RNA binding site of MSI2 RRM1. Distances in Å; (f) Docked pose of Ro-NGF in the RNA binding site of MSI2 RRM1 showing a displaced center of the small-molecule from the binding site; (g) 2D representation of Ro-OH docked pose in the RRM1 of MSI2; (h) 2D representation of Ro-NGF docked pose in the RRM1 of MSI2 showing H-bonding of K22 changing from the O to the N in the middle ring, and π-cation interacting with R100 displaced with respect to Ro (see ***Figure 2b***).

**Extended Data Figure 3.**
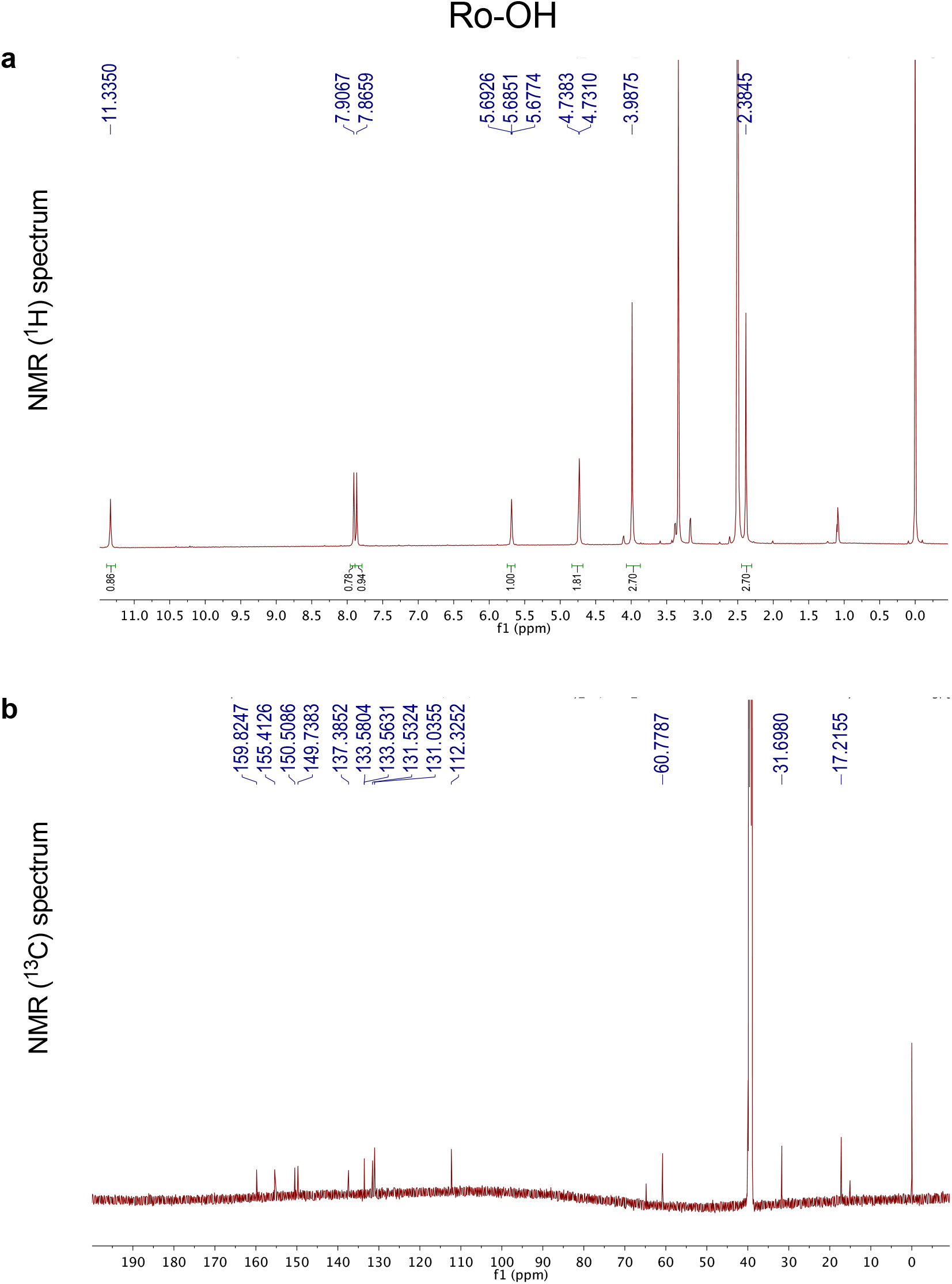
^1^**H NMR** (a) **and NMR^13^C spectrum** (b) **of Ro-OH, the synthesized reduced form of Ro**.

**Extended Data Figure 4.**
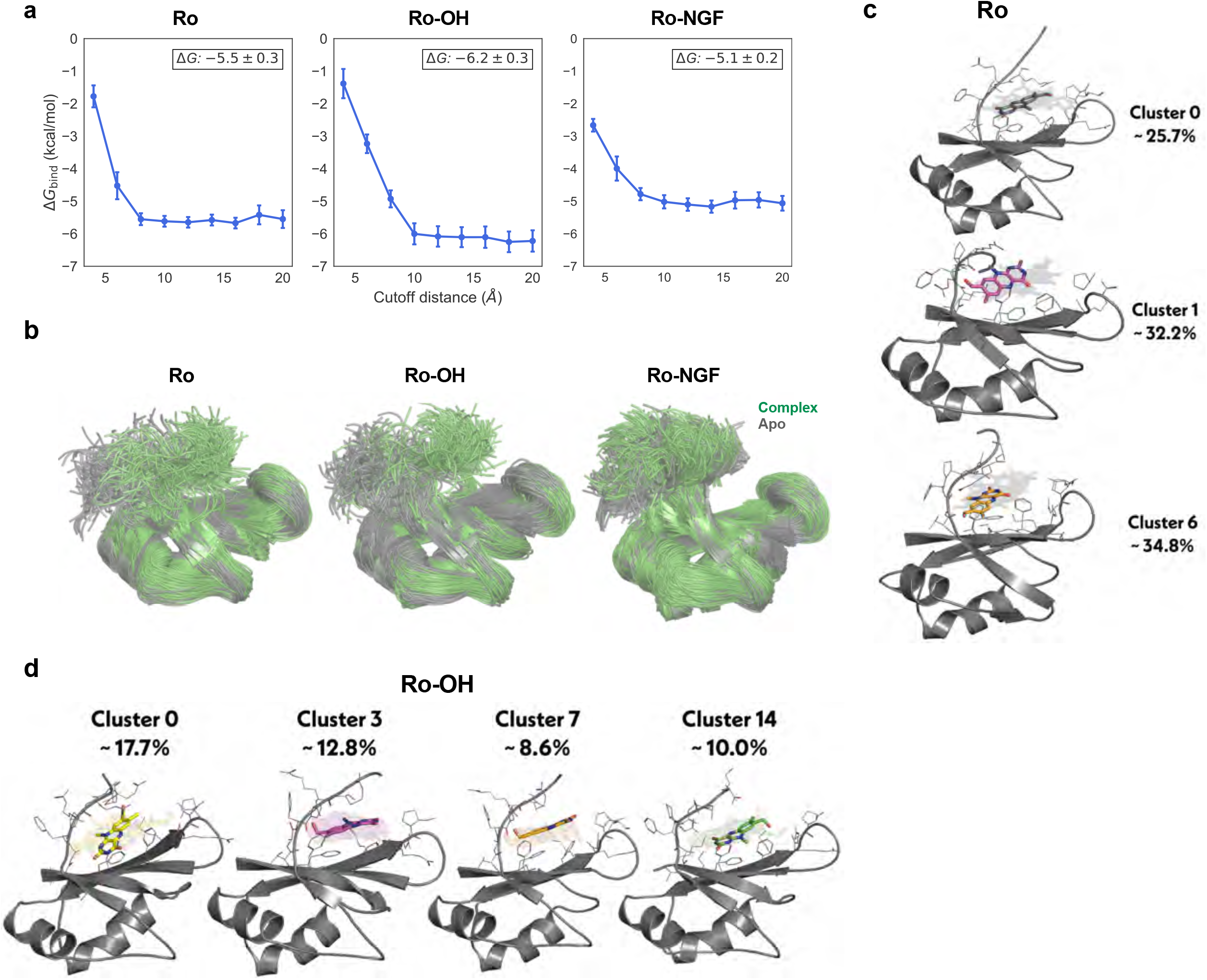
Alchemical free energy calculations show that both protein and ligands adopt a conformationally heterogeneous ensemble of binding poses. (a) Computed binding free energy (ΔG_bind_, kcal/mol) estimates from alchemical free energy calculations (*y-axis*) for Ro, Ro-OH, and Ro-NGF for different definitions of the “bound” complex as a function of distance cutoff (*x-axis*, in Å). Reported statistical errors and error bars correspond one standard error. The inset ΔG_bind_ was calculated for a cutoff of 20Å. (b) In the alchemical Hamiltonian replica exchange simulations, a conformational change is induced when MSI2 is bound (“Complex”; *green*) to Ro (*right*) or Ro-OH (*center*), as compared to apo MSI2 (“Apo”; *gray*). Ro-NGF (*left*) does not induce the same conformational change. (c) The top three most populous clusters for Ro 08–2750. The protein structure and solid-color ligand pose depict cluster centers, while transparent ligand poses depict 10 randomly sampled frames assigned to that cluster. Sidechains within 4Å of any of the ligands are shown as lines. (d) The top four most populous clusters for Ro-OH, using the same depiction scheme as (c).

**Extended Data Figure 5.**
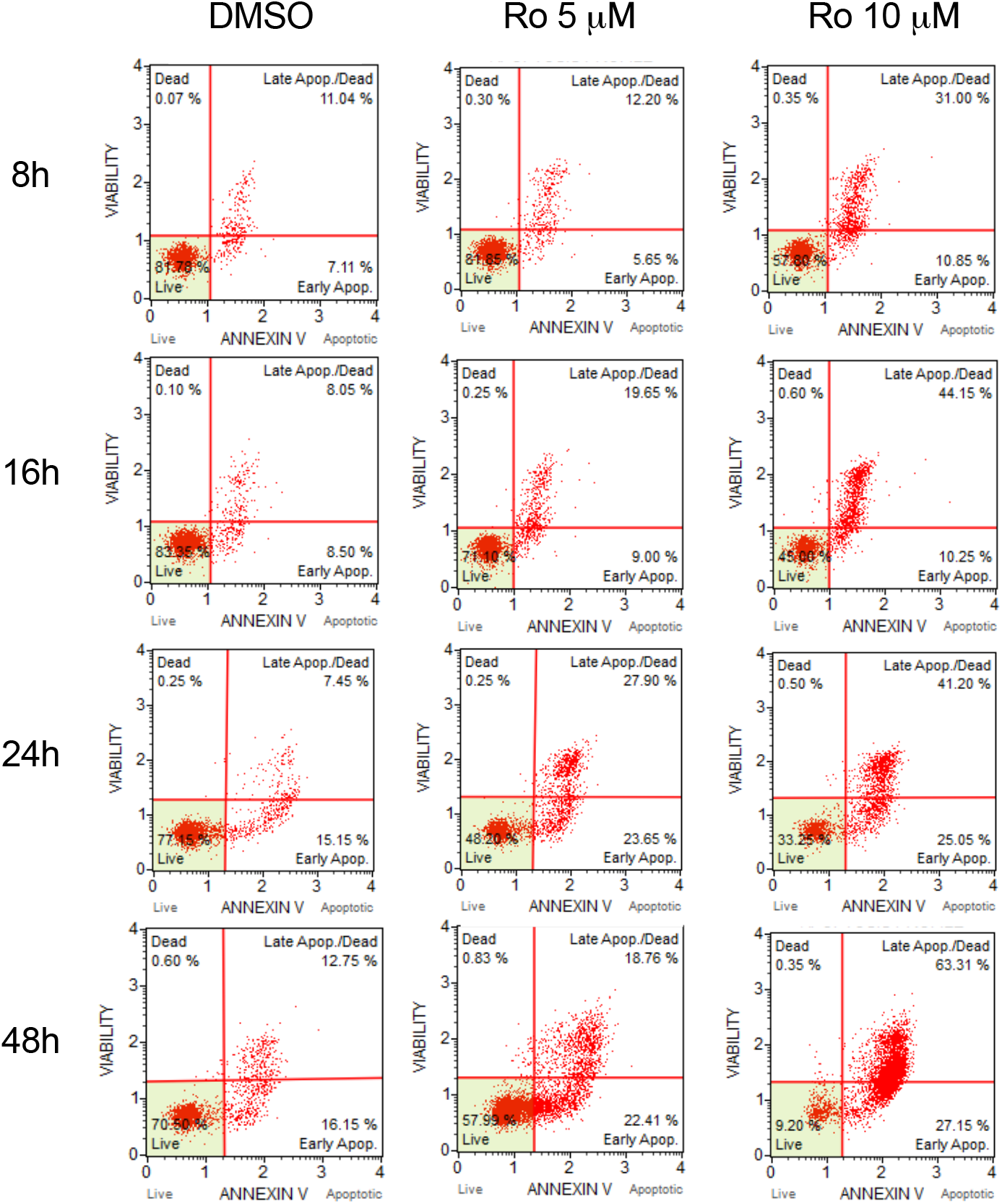
Flow cytometry plots showing apoptosis in MLL-AF9 leukemic cells after treatment with Ro 08–2750. Apoptosis plots (graphs in Figure 3d) showing Annexin V+ and 7AAD (live/dead staining) by Apoptosis MUSE^®^ Cell kit and MUSE^®^ Cell Analyzer (Millipore-Sigma) in MLL-AF9+ BM cells at 8, 16, 24 and 48 hours post treatment with Ro 5 and 10 μM.

**Extended Data Figure 6.**
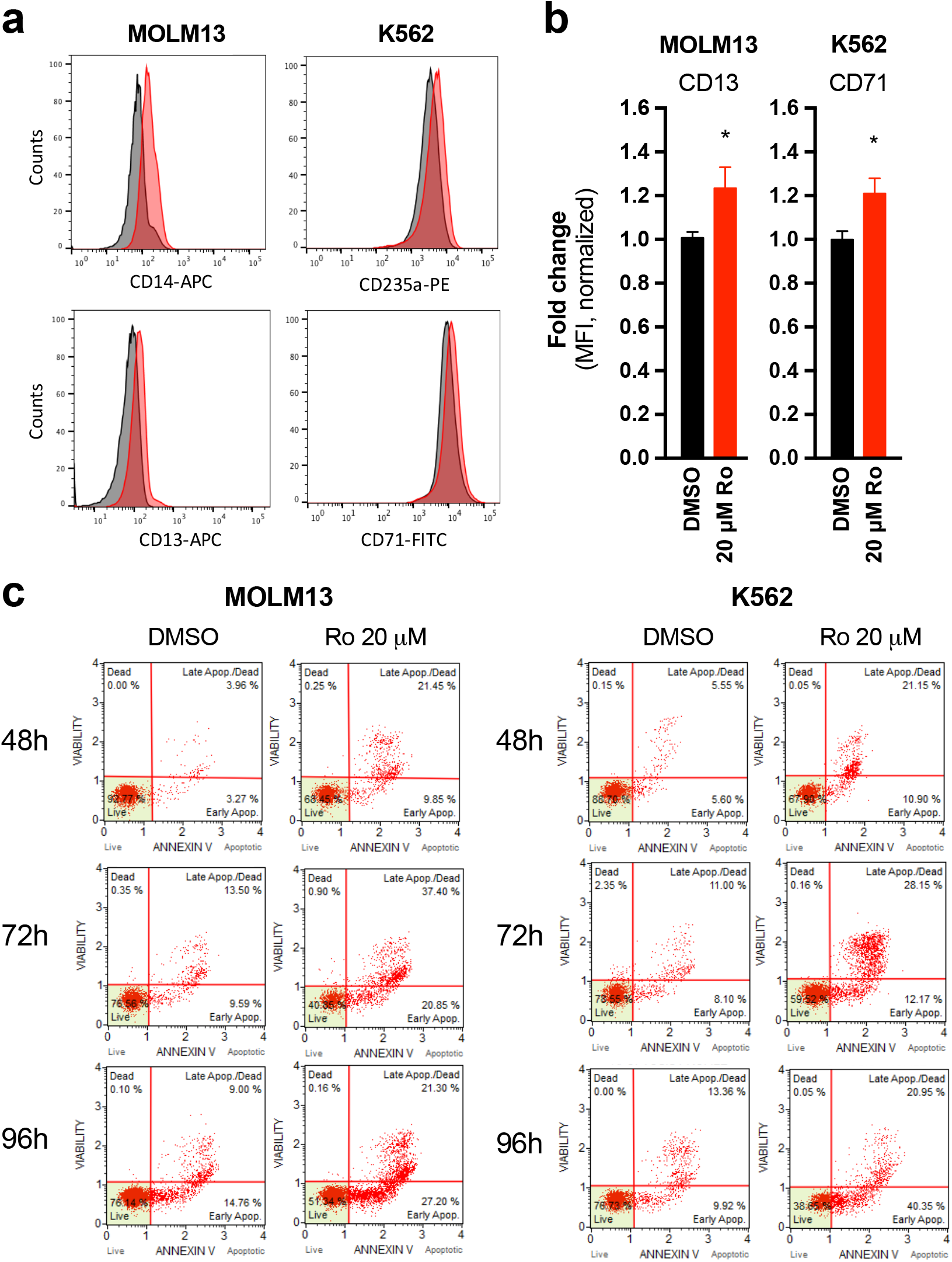
Differentiation and apoptosis are induced in MOLM13 and K562 cells after Ro 08–2750 treatment. (a) Representative histograms showing CD14 and CD13 myeloid markers in MOLM13 and erythroid differentiation markers CD235a (Glycophorin-A) and CD71 in K562 after 48h of 20 μM Ro treatment. (b) Mean Fluorescence Intensity (MFI) fold changes of CD13 (myeloid marker, MOLM13) and CD71 (erythroid marker, K562) after 48h treatment of leukemia cell lines with DMSO (control, *black bars*) or Ro 20 μM (*red bars*). Data is shown as average (normalized to DMSO control cells) ± standard error mean of three independent experiments performed in triplicate. Paired *t*-test (DMSO vs Ro treated); \**p*<0.05. (c) Apoptosis plots (from graphs in Figure 4d) showing Annexin V+ and 7AAD (live/dead staining) in MOLM13 and K562 by MUSE^®^ Cell Analyzer (Millipore-Sigma) in DMSO and Ro 20 μM treatments at 48, 72 and 96h.

**Extended Data Figure 7.**
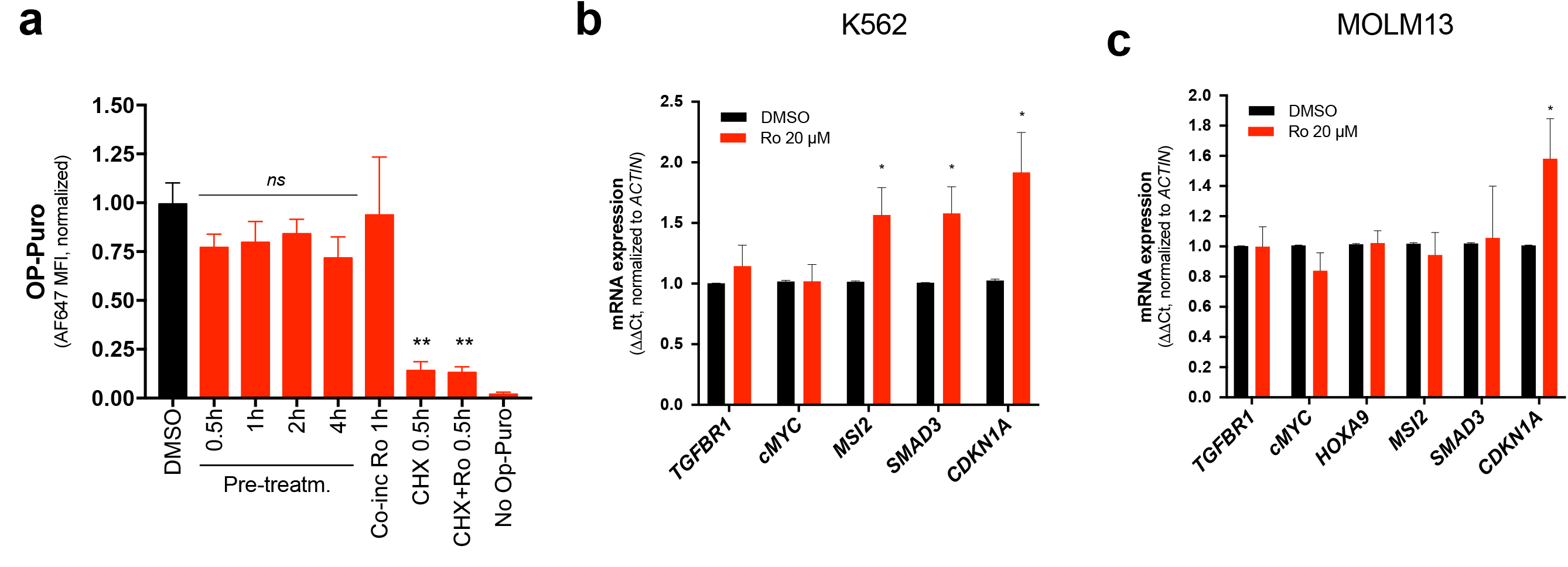
Ro effects on global translation and mRNA of MSI2 targets. (a) OP-Puromycin incorporation to assess global translation rates in MOLM13 leukemia cells. Results are represented as average of Alexa Fluor 647 (AF647) Mean Fluorescence Intensity (MFI) normalized to DMSO control cells ± standard error mean of four independent experiments performed in duplicate. Paired *t*-test (DMSO vs Ro treated); *ns*, non-significant, \*\**p*<0.005. (b) Expression levels of mRNA targets of MSI2 by qPCR in K562 and (c) MOLM13. Cells were treated for 4h at 20 μM Ro. Results represent the average of ten independent experiments ± standard error mean. Paired *t*-test (DMSO vs Ro treated); \**p*<0.05.

**Extended Data Figure 8.**
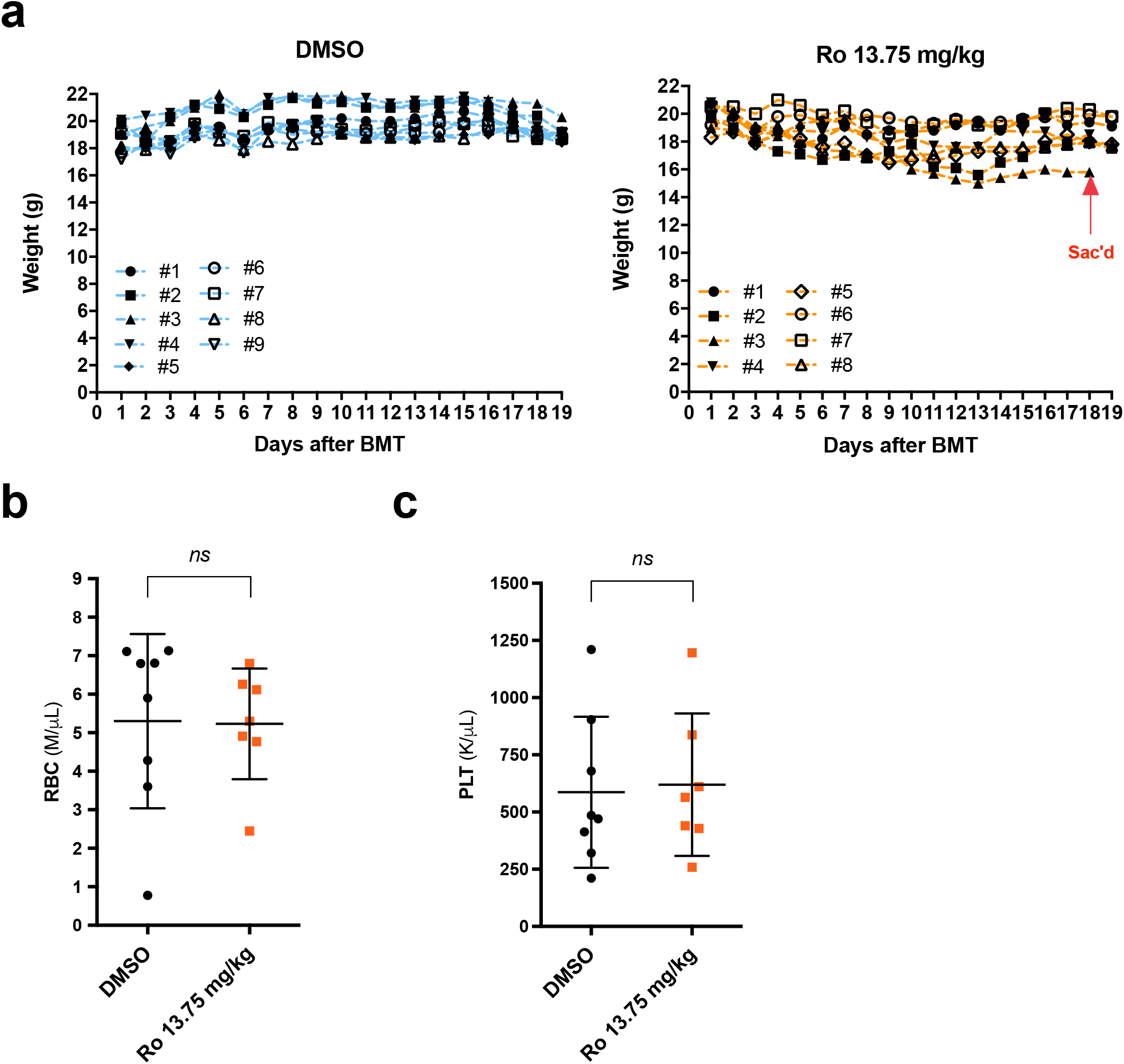
No toxicity of Ro 08–2750 after in vivo treatment of MLL-AF9 mice. (a) Mice weight in DMSO (*cyan lines*, left panel) and Ro 13.75 mg/kg (*orange lines*, right panel) groups during the duration of the *in vivo* experiment. (b) Red Blood Cell (RBC) counts (M/ μL) at time of sacrifice. Each data point represents an individually treated mouse. Unpaired *t*-test; *ns*, non-significant. (c) Platelets counts (PLT) counts (K/ μL) at time of sacrifice. Each data point represents an individually treated mouse. Unpaired *t*-test; *ns*, non-significant. DMSO, *n* = 9; Ro, *n* = 8.

## Acknowledgements

We would like to thank the current members of the M. Kharas and R. Levine labs, R. Levine, P. Lito, J. Vidigal, P. Rocha, C. Lengner, A. Kentsis and K. Keshari for their critical advice and helpful suggestions. H. Djaballah, R. Garippa and the members of the former High-Throughput Screening Core and current RNAi Core (MSKCC) for their technical and project support throughout the screen and secondary validation. We would like to thank R. Sridharan, Y. Shamay, D. Heller (MSKCC) and Rui Liang (Tri-TDI) for their scientific advice and technical support. We would also like to thank A. Gruet and the Epigenetics core for his technical support with RNA-sequencing library preparation and J. Charavalli (HTSRC, The Rockefeller University) for her technical support with initial FP measurements. M.G.K. was supported by US National Institutes of Health National Institute of Diabetes, Digestive and Kidney Diseases Career Development Award, NIDDK NIH R01-DK101989–01A1, NCI 1R01CA193842–01, Louis V. Gerstner Young Investigator Award, American Society of Hematology Junior Scholar Award, Kimmel Scholar Award, V-Scholar Award, Geoffrey Beene Award and Alex’s Lemonade Stand A Award and the Starr Foundation. The research was funded by Mr. William H. Goodwin and Mrs. Alice Goodwin and the Commonwealth Foundation for Cancer Research and The Center for Experimental Therapeutics at Memorial Sloan Kettering Cancer Center. The research was funded in part through NIH/NCI Cancer Support Core Grant P30 CA08748 to M.G.K. J.D.C. acknowledges support from the Sloan Kettering Institute, a Louis V. Gerstner Young Investigator Award, NIH grant P30 CA008748, and NIH grant R01 GM121505. L.N.N. acknowledges support from Merck KGaA to support the development of open source tools for GPU-accelerated alchemical free energy calculations. M.C.P was supported by a Medical Scientist Training Program grant from the National Institute of General Medical Sciences of the National Institutes of Health under award number T32GM007739 to the Weill Cornell/Rockefeller/Sloan Kettering Tri-Institutional MD-PhD Program. The MSKCC structural biology core laboratory is supported by National Cancer Institute grant P30-CA008748. X-ray diffraction data were collected at synchrotron facilities supported by grants and contracts from the National Institutes of Health (P41GM103403, HEI-S10RR029205) and the Department of Energy (DE-AC02–06CH11357).

## Author Contributions

G.M. led the project, performed experiments, analyzed data and wrote the manuscript. M.G.K. directed the project, analyzed data and wrote the manuscript. S.A., D.C., T.B., A.R., L.N., J.C. performed experiments, analyzed data and provided project support. A.C., A.S., S.M.P., T.C., J.T. performed experiments and analyzed data. M.C.P., L.F., C.L. analyzed data. J.S., C.F., M.P. provided clinical data and analysis. C.Z.R. and D.T. performed experiments and provided critical reagents. J.E. and G.M.R. provided critical reagents. Y.G. performed experiments and analyzed data. C.A. and J.F.G. provided suggestions, project support and assisted analyzing data.

## Author Information

J.D.C. is a member of the Scientific Advisory Board for Schrödinger.

## Methods. Supplemental References

1 Park, S. M. et al. Musashi2 sustains the mixed-lineage leukemia-driven stem cell regulatory program. J Clin Invest 125, 1286–1298 (2015).

2 Minuesa, G. et al. A 1536-well fluorescence polarization assay to screen for modulators of the MUSASHI family of RNA-binding proteins. Comb Chem High Throughput Screen 17, 596–609 (2014).

3 Otwinowski, Z. & Minor, W. Processing of X-ray diffraction data collected in oscillation mode. Methods Enzymol 276, 307–326 (1997).

4 Adams, P. D. et al. PHENIX: a comprehensive Python-based system for macromolecular structure solution. Acta Crystallogr D Biol Crystallogr 66, 213–221 (2010).

5 Jones, T. A., Zou, J. Y., Cowan, S. W. & Kjeldgaard, M. Improved methods for building protein models in electron density maps and the location of errors in these models. Acta Crystallogr A 47 (Pt 2), 110–119 (1991).

6 Salach, J. et al. Studies on succinate dehydrogenase. Site of attachment of the covalently-bound flavin to the peptide chain. Eur J Biochem 26, 267–278 (1972).

7 Sastry, G. M., Adzhigirey, M., Day, T., Annabhimoju, R. & Sherman, W. Protein and ligand preparation: parameters, protocols, and influence on virtual screening enrichments. J Comput Aided Mol Des 27, 221–234 (2013).

8 Halgren, T. A. Identifying and characterizing binding sites and assessing druggability. J Chem Inf Model 49, 377–389 (2009).

9 Halgren, T. New method for fast and accurate binding-site identification and analysis. Chem Biol Drug Des 69, 146–148 (2007).

10 Halgren, T. A. et al. Glide: a new approach for rapid, accurate docking and scoring. 2. Enrichment factors in database screening. J Med Chem 47, 1750–1759 (2004).

11 Friesner, R. A. et al. Glide: a new approach for rapid, accurate docking and scoring. 1. Method and assessment of docking accuracy. J Med Chem 47, 1739–1749 (2004).

12 Lan, L. et al. Natural product (-)-gossypol inhibits colon cancer cell growth by targeting RNA-binding protein Musashi-1. Mol Oncol 9, 1406–1420 (2015).

13 Sherman, W., Day, T., Jacobson, M. P., Friesner, R. A. & Farid, R. Novel procedure for modeling ligand/receptor induced fit effects. J Med Chem 49, 534–553 (2006).

14 Clark, A. J. et al. Prediction of Protein-Ligand Binding Poses via a Combination of Induced Fit Docking and Metadynamics Simulations. J Chem Theory Comput 12, 2990–2998 (2016).

15 Young, T., Abel, R., Kim, B., Berne, B. J. & Friesner, R. A. Motifs for molecular recognition exploiting hydrophobic enclosure in protein-ligand binding. Proc Natl Acad Sci U S A 104, 808–813 (2007).

16 Abel, R., Young, T., Farid, R., Berne, B. J. & Friesner, R. A. Role of the active-site solvent in the thermodynamics of factor Xa ligand binding. J Am Chem Soc 130, 2817–2831, (2008).

17 Case, D. A., Betz, R.M., Cerutti, D.S., Cheatham, III, T.E., Darden, T.A., Duke, R.E., Giese, T.J., Gohlke, H., Goetz, A.W., Homeyer, N., Izadi, S., Janowski, P., Kaus, J., Kovalenko, A., Lee, T.S., LeGrand, S., Li, P., Lin, C., Luchko, T., Luo, R., Madej, R., Mermelstein, D., Merz, K.M., Monard, G., Nguyen, H., Nguyen, H.T., Omelyan, I., Onufriev, A., Roe, D.R., Roitberg, A., Sagui, C., Simmerling, C.L., Botello-Smith, W.M., Swails, J., Walker, R.C., Wang, J., Wolf, R.M., Wu, X., Xiao, L. and Kollman, P.A. AMBER 2016, University of California, San Francisco (2016).

18 Maier, J. A. et al. ff14SB: Improving the Accuracy of Protein Side Chain and Backbone Parameters from ff99SB. J Chem Theory Comput 11, 3696–3713 (2015).

19 Wang, J., Wolf, R. M., Caldwell, J. W., Kollman, P. A. & Case, D. A. Development and testing of a general amber force field. J Comput Chem 25, 1157–1174 (2004).

20 Wang, J., Wang, W., Kollman, P. A. & Case, D. A. Automatic atom type and bond type perception in molecular mechanical calculations. J Mol Graph Model 25, 247–260 (2006).

21 Jakalian, A., Jack, D. B. & Bayly, C. I. Fast, efficient generation of high-quality atomic charges. AM1-BCC model: II. Parameterization and validation. J Comput Chem 23, 1623–1641 (2002).

22 Jakalian, A., Bush, B. L., Jack, D. B. & Bayly, C. I. Fast, efficient generation of high-quality atomic carges AM1-BCC model: I. Method. J Comput Chem 21, 132–146 (2000).

23 Toolkits, O. Oct.1 OpenEye Scientific Software, Santa Fe, NM. http://www.eyesopen.com/. (2017).

24 Nocedal, J. American Mathematical Society. Math. Comp. 35, 773–782 (1980).

25 Eastman, P. et al. OpenMM 7: Rapid development of high performance algorithms for molecular dynamics. PLoS Comput Biol 13, E1005659f (2017).

26 Wang, K., Chodera, J. D., Yang, Y. & Shirts, M. R. Identifying ligand binding sites and poses using GPU-accelerated Hamiltonian replica exchange molecular dynamics. J Comput Aided Mol Des 27, 989–1007 (2013).

27 Darden, T., York, D. & Pedersen, L. Particle mesh Ewald: An N log (N) method for Ewald sums in large systems J Chem Phys 98, 10089–10092 (1998).

28 Jorgensen, W. L., Chandrasekhar, J., Madura, J. D., Impey, R. W. & Klein, M. L. Comparison of simple potential functions for simulating liquid water. J Chem Phys 79, 926–935 (1998).

29 Joung, I. S. & Cheatham, T. E., 3rd. Determination of alkali and halide monovalent ion parameters for use in explicitly solvated biomolecular simulations. J Phys Chem B 112, 9020–9041 (2008).

30 Leimkuhler, B. & Matthews, C. Efficient molecular dynamics using geodesic integration and solvent-solute splitting. Proc Math Phys Eng Sci 472, 20160138 (2016).

31 Chodera, J. D. & Shirts, M. R. Replica exchange and expanded ensemble simulations as Gibbs sampling: simple improvements for enhanced mixing. J Chem Phys 135, 194110 (2011).

32 Shirts, M. R. & Chodera, J. D. Statistically optimal analysis of samples from multiple equilibrium states. J Chem Phys 129, 124105 (2008).

33 Chodera, J. D. A Simple Method for Automated Equilibration Detection in Molecular Simulations. J Chem Theory Comput 12, 1799–1805 (2016).

34 McGibbon, R. T. et al. MDTraj: A Modern Open Library for the Analysis of Molecular Dynamics Trajectories. Biophys J 109, 1528–1532 (2015).

35 Beauchamp, K. A. et al. MSMBuilder2: Modeling Conformational Dynamics at the Picosecond to Millisecond Scale. J Chem Theory Comput 7, 3412–3419 (2011).

## References

1 Hentze, M. W., Castello, A., Schwarzl, T. & Preiss, T. A brave new world of RNA-binding proteins. Nat Rev Mol Cell Biol, doi:10.1038/nrm.2017.130 (2018).

2 Kharas, M. G. & Lengner, C. J. Stem Cells, Cancer, and MUSASHI in Blood and Guts. Trends Cancer 3, 347–356, doi:10.1016/j.trecan.2017.03.007 (2017).

3 Pereira, B., Billaud, M. & Almeida, R. RNA-Binding Proteins in Cancer: Old Players and New Actors. Trends Cancer 3, 506–528, doi:10.1016/j.trecan.2017.05.003 (2017).

4 Vu, L. P. et al. Functional screen of MSI2 interactors identifies an essential role for SYNCRIP in myeloid leukemia stem cells. Nat Genet 49, 866–875, doi:10.1038/ng.3854 (2017).

5 Han, T. et al. Anticancer sulfonamides target splicing by inducing RBM39 degradation via recruitment to DCAF15. Science 356, doi:10.1126/science.aal3755 (2017).

6 Ghosh, M. et al. Essential role of the RNA-binding protein HuR in progenitor cell survival in mice. J Clin Invest 119, 3530–3543, doi:10.1172/JCI38263 (2009).

7 Palanichamy, J. K. et al. RNA-binding protein IGF2BP3 targeting of oncogenic transcripts promotes hematopoietic progenitor proliferation. J Clin Invest 126, 1495–1511, doi:10.1172/JCI80046 (2016).

8 Park, S. M. et al. Musashi2 sustains the mixed-lineage leukemia-driven stem cell regulatory program. J Clin Invest 125, 1286–1298, doi:10.1172/JCI78440 (2015).

9 Lee, S. C. & Abdel-Wahab, O. Therapeutic targeting of splicing in cancer. Nat Med 22, 976–986, doi:10.1038/nm.4165 (2016).

10 Kanemura, Y. et al. Musashi1, an evolutionarily conserved neural RNA-binding protein, is a versatile marker of human glioma cells in determining their cellular origin, malignancy, and proliferative activity. Differentiation 68, 141–152 (2001).

11 Hemmati, H. D. et al. Cancerous stem cells can arise from pediatric brain tumors. Proc Natl Acad Sci U S A 100, 15178–15183, doi:10.1073/pnas.2036535100 (2003).

12 Shu, H. J. et al. Expression of the Musashi1 gene encoding the RNA-binding protein in human hepatoma cell lines. Biochem Biophys Res Commun 293, 150–154, doi:10.1016/S0006-291X(02)00175-4 (2002).

13 Li, N. et al. The Msi Family of RNA-Binding Proteins Function Redundantly as Intestinal Oncoproteins. Cell Rep 13, 2440–2455, doi:10.1016/j.celrep.2015.11.022 (2015).

14 Wang, S. et al. Transformation of the intestinal epithelium by the MSI2 RNA-binding protein. Nat Commun 6, 6517, doi:10.1038/ncomms7517 (2015).

15 Oskarsson, T. et al. Breast cancer cells produce tenascin C as a metastatic niche component to colonize the lungs. Nat Med 17, 867–874, doi:10.1038/nm.2379 (2011).

16 Kang, M. H. et al. Musashi RNA-binding protein 2 regulates estrogen receptor 1 function in breast cancer. Oncogene 36, 1745–1752, doi:10.1038/onc.2016.327 (2017).

17 Wang, X. Y. et al. Musashi1 as a potential therapeutic target and diagnostic marker for lung cancer. Oncotarget 4, 739–750, doi:10.18632/oncotarget.1034 (2013).

18 Vo, D. T. et al. The oncogenic RNA-binding protein Musashi1 is regulated by HuR via mRNA translation and stability in glioblastoma cells. Mol Cancer Res 10, 143–155, doi:10.1158/1541-7786.MCR-11-0208 (2012).

19 Guo, K. et al. The Novel KLF4/MSI2 Signaling Pathway Regulates Growth and Metastasis of Pancreatic Cancer. Clin Cancer Res 23, 687–696, doi:10.1158/1078-0432.CCR-16-1064 (2017).

20 Fox, R. G. et al. Image-based detection and targeting of therapy resistance in pancreatic adenocarcinoma. Nature 534, 407–411, doi:10.1038/nature17988 (2016).

21 Barbouti, A. et al. A novel gene, MSI2, encoding a putative RNA-binding protein is recurrently rearranged at disease progression of chronic myeloid leukemia and forms a fusion gene with HOXA9 as a result of the cryptic t (7;17)(p15;q23). Cancer Res 63, 1202–1206 (2003).

22 De Weer, A. et al. EVI1 overexpression in t (3;17) positive myeloid malignancies results from juxtaposition of EVI1 to the MSI2 locus at 17q22. Haematologica 93, 1903–1907, doi:10.3324/haematol.13192 (2008).

23 Saleki, R. et al. A novel TTC40-MSI2 fusion in de novo acute myeloid leukemia with an unbalanced 10;17 translocation. Leuk Lymphoma 56, 1137–1139, doi:10.3109/10428194.2014.947611 (2015).

24 Wang, K. et al. Patient-derived xenotransplants can recapitulate the genetic driver landscape of acute leukemias. Leukemia 31, 151–158, doi:10.1038/leu.2016.166 (2017).

25 Ito, T. et al. Regulation of myeloid leukaemia by the cell-fate determinant Musashi. Nature 466, 765–768, doi:10.1038/nature09171 (2010).

26 Kharas, M. G. et al. Musashi-2 regulates normal hematopoiesis and promotes aggressive myeloid leukemia. Nat Med 16, 903–908, doi:10.1038/nm.2187 (2010).

27 Thol, F. et al. Prognostic significance of expression levels of stem cell regulators MSI2 and NUMB in acute myeloid leukemia. Ann Hematol 92, 315–323, doi:10.1007/s00277-012-1637-5 (2013).

28 Taggart, J. et al. MSI2 is required for maintaining activated myelodysplastic syndrome stem cells. Nat Commun 7, 10739, doi:10.1038/ncomms10739 (2016).

29 Byers, R. J., Currie, T., Tholouli, E., Rodig, S. J. & Kutok, J. L. MSI2 protein expression predicts unfavorable outcome in acute myeloid leukemia. Blood 118, 2857–2867, doi:10.1182/blood-2011-04-346767 (2011).

30 Kwon, H. Y. et al. Tetraspanin 3 Is Required for the Development and Propagation of Acute Myelogenous Leukemia. Cell Stem Cell 17, 152–164, doi:10.1016/j.stem.2015.06.006 (2015).

31 Park, S. M. et al. Musashi-2 controls cell fate, lineage bias, and TGF-beta signaling in HSCs. J Exp Med 211, 71–87, doi:10.1084/jem.20130736 (2014).

32 Rentas, S. et al. Musashi-2 attenuates AHR signalling to expand human haematopoietic stem cells. Nature 532, 508–511, doi:10.1038/nature17665 (2016).

33 Kudinov, A. E., Karanicolas, J., Golemis, E. A. & Boumber, Y. Musashi RNA-Binding Proteins as Cancer Drivers and Novel Therapeutic Targets. Clin Cancer Res 23, 2143–2153, doi:10.1158/1078-0432.CCR-16-2728 (2017).

34 Sakakibara, S., Nakamura, Y., Satoh, H. & Okano, H. Rna-binding protein Musashi2: developmentally regulated expression in neural precursor cells and subpopulations of neurons in mammalian CNS. J Neurosci 21, 8091–8107 (2001).

35 Zearfoss, N. R. et al. A conserved three-nucleotide core motif defines Musashi RNA binding specificity. J Biol Chem 289, 35530–35541, doi:10.1074/jbc.M114.597112 (2014).

36 Ohyama, T. et al. Structure of Musashi1 in a complex with target RNA: the role of aromatic stacking interactions. Nucleic Acids Res 40, 3218–3231, doi:10.1093/nar/gkr1139 (2012).

37 Katz, Y. et al. Musashi proteins are post-transcriptional regulators of the epithelial-luminal cell state. Elife 3, e03915, doi:10.7554/eLife.03915 (2014).

38 Minuesa, G. et al. A 1536-well fluorescence polarization assay to screen for modulators of the MUSASHI family of RNA-binding proteins. Comb Chem High Throughput Screen 17, 596–609 (2014).

39 Eibl, J. K., Strasser, B. C. & Ross, G. M. Identification of novel pyrazoloquinazolinecarboxilate analogues to inhibit nerve growth factor in vitro. Eur J Pharmacol 708, 30–37, doi:10.1016/j.ejphar.2013.03.029 (2013).

40 Subramanian, A. et al. Gene set enrichment analysis: a knowledge-based approach for interpreting genome-wide expression profiles. Proc Natl Acad Sci U S A 102, 15545–15550, doi:10.1073/pnas.0506580102 (2005).

41 Zhang, H. et al. Musashi2 modulates K562 leukemic cell proliferation and apoptosis involving the MAPK pathway. Exp Cell Res 320, 119–127, doi:10.1016/j.yexcr.2013.09.009 (2014).

42 Han, Y. et al. Musashi-2 Silencing Exerts Potent Activity against Acute Myeloid Leukemia and Enhances Chemosensitivity to Daunorubicin. PLoS One 10, e0136484, doi:10.1371/journal.pone.0136484 (2015).

43 Meisner, N. C. et al. Identification and mechanistic characterization of low-molecular-weight inhibitors for HuR. Nat Chem Biol 3, 508–515, doi:10.1038/nchembio.2007.14 (2007).

44 Wu, X. et al. Identification and validation of novel small molecule disruptors of HuRmRNA interaction. ACS Chem Biol 10, 1476–1484, doi:10.1021/cb500851u (2015).

45 Lan, L. et al. Natural product (-)-gossypol inhibits colon cancer cell growth by targeting RNA-binding protein Musashi-1. Mol Oncol 9, 1406–1420, doi:10.1016/j.molonc.2015.03.014 (2015).

46 Lim, D., Byun, W. G., Koo, J. Y., Park, H. & Park, S. B. Discovery of a Small-Molecule Inhibitor of Protein-MicroRNA Interaction Using Binding Assay with a Site-Specifically Labeled Lin28. J Am Chem Soc, doi:10.1021/jacs.6b06965 (2016).

47 Roos, M. et al. A Small-Molecule Inhibitor of Lin28. ACS Chem Biol 11, 2773–2781, doi:10.1021/acschembio.6b00232 (2016).

48 Jarvis, W. D., Turner, A. J., Povirk, L. F., Traylor, R. S. & Grant, S. Induction of apoptotic DNA fragmentation and cell death in HL-60 human promyelocytic leukemia cells by pharmacological inhibitors of protein kinase C. Cancer Res 54, 1707–1714 (1994).

49 Zhu, J. et al. Niemann-Pick C2 Proteins: A New Function for an Old Family. Front Physiol 9, 52, doi:10.3389/fphys.2018.00052 (2018).

50 Judge, J. L. et al. The Lactate Dehydrogenase Inhibitor Gossypol Inhibits Radiation-Induced Pulmonary Fibrosis. Radiat Res 188, 35–43, doi:10.1667/RR14620.1 (2017).

51 Zeng, Y., Ma, J., Xu, L. & Wu, D. Natural Product Gossypol and Its Derivatives in Precision Cancer Medicine. Curr Med Chem, doi:10.2174/0929867324666170523123655 (2017).

52 Clingman, C. C. et al. Allosteric inhibition of a stem cell RNA-binding protein by an intermediary metabolite. Elife 3, doi:10.7554/eLife.02848 (2014).

53 Sadlish, H. et al. Evidence for a functionally relevant rocaglamide binding site on the eIF4A-RNA complex. ACS Chem Biol 8, 1519–1527, doi:10.1021/cb400158t (2013).

54 Choo, A. Y., Yoon, S. O., Kim, S. G., Roux, P. P. & Blenis, J. Rapamycin differentially inhibits S6Ks and 4E-BP1 to mediate cell-type-specific repression of mRNA translation. Proc Natl Acad Sci U S A 105, 17414–17419, doi:10.1073/pnas.0809136105 (2008).

55 Fang, T. et al. Musashi 2 contributes to the stemness and chemoresistance of liver cancer stem cells via LIN28A activation. Cancer Lett 384, 50–59, doi:10.1016/j.canlet.2016.10.007 (2017).

56 Sheng, W. et al. Cooperation of Musashi-2, Numb, MDM2, and P53 in drug resistance and malignant biology of pancreatic cancer. FASEB J 31, 2429–2438, doi:10.1096/fj.201601240R (2017).

